# One species, two lives: cryptic population structure and life-history divergence in a vineyard spittlebug (*Aphrophora sp.*), a candidate vector for Pierce’s disease

**DOI:** 10.64898/2026.09.21.753153

**Authors:** Manpreet K. Kohli, Ethan R. Tolman, Vinton Thompson

**Affiliations:** Department of Natural Sciences, Baruch College, City University of New York, New York, USA; CUNY Graduate Center, New York, NY, USA; Division of Invertebrate Zoology, American Museum of Natural History, New York, USA; Department of Biological Sciences, Virginia Tech, Virginia, USA

**Keywords:** Biogeography, population genomics, Cercopoidea, Aphrophoridae, *Xylella fastidiosa*

## Abstract

Whole genome analysis of spittlebug populations, an *Aphrophora* species (Hemiptera: Aphrophoridae) known to spread Pierce’s disease, from various vineyards in Napa and Sonoma Counties demonstrates that these insects are conspecific with *Aphrophora permutata* found in quasi-natural populations. The genomes analyzed fall in three autosomal clusters with two sympatric populations (based on admixture) that diverged around 40,000-90,000 years before present according to demographic models. We also recover three related but distinct mitochondrial clusters. There is no genetic differentiation between Napa and Sonoma County vineyard populations and very modest differentiation among vineyards and quasi-natural Berkeley/Oakland Hills population. These vineyard and quasi-natural populations have distinct life histories, including divergent oviposition hosts (understory herbs versus pines) and different patterns of adult seasonal migration (out-migration and return versus simple movement from the understory to the pine canopy). Analysis of the mitochondrial gene COI indicates that *A. permutata*, Aphrophora fulva and *Aphrophora maculosa* are intermingled in the same mitochondrial clade, along with the more distantly related species *Aphrophora gelida*. Further, all three autosomal genetic clusters show evidence of selection for insecticide resistance. We also find that endosymbionts *Wolbachia* and *Rickettsia* were both present in a substantial fraction of specimens analyzed, but neither appears to influence population structure. Overall, our findings elucidate an intriguing degree of life history divergence between geographically proximate, conspecific populations linked by gene flow and may have implications for the continued management of vineyard populations.

## Introduction

Land-use change is a significant driver of biodiversity decline (Maxwell et al., 2016). Specifically, agricultural expansion and intensification affects 62% of assessed species classified as threatened or near-threatened by IUCN, while urban development has negatively impacted 35% of such species (Maxwell et al., 2016). Against this backdrop of biodiversity decline and highly altered landscapes, a small handful of species have been able to thrive. Such species, the so-called “winners” of the Anthropocene, present major challenges to human health and economic activity. Agricultural arthropod pests especially present a major threat to global food security, destroying 18-20% of crop production globally (Sharma et al., 2017).

Such pests include some spittlebugs (Hemiptera: Cercopoidea) whose nymphs develop within characteristic protective foam masses (Fig. 1C) and are xylem-sap feeders. Spittlebugs can inflict direct feeding damage and play a major role in transmitting *Xylella fastidiosa*, a bacterium that causes Pierce’s disease of grapes, olive quick decline syndrome, and several other diseases in crops (Thompson, 2026). The meadow spittlebug, *Philaenus spumarius* (Aphrophoridae), one of the world’s most widespread and abundant insects, is the best-studied example, transmitting *X. fastidiosa* to hosts as diverse as grapes, almonds, and citrus (Cornara et al., 2016, 2017). With an extraordinary feeding breadth (631 plant genera and 117 families) *P. spumarius* is primed to spread *X. fastidiosa* further (Thompson et al., 2023).

**Fig. 1.**
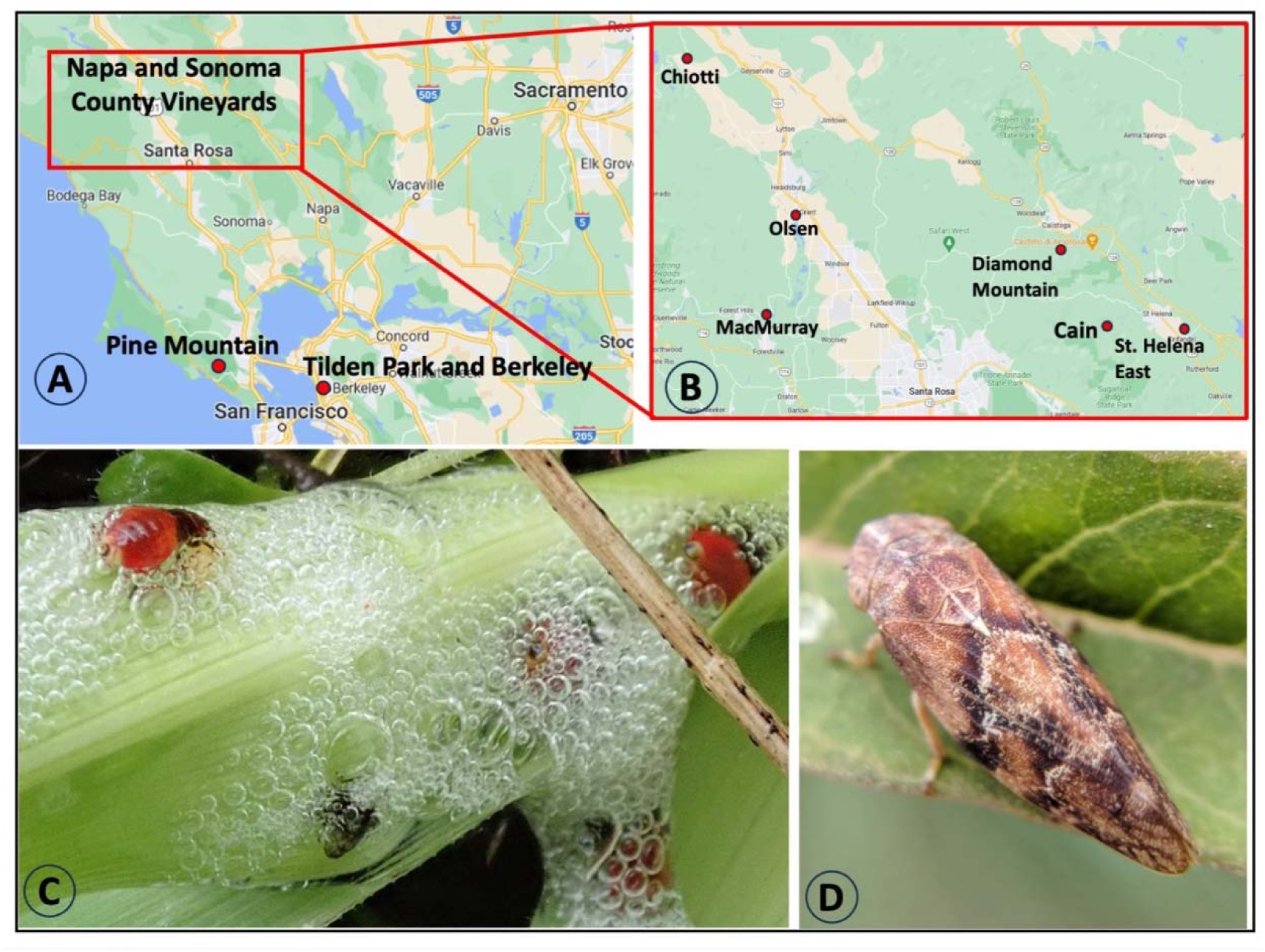
Study Design. A. and B. Map of California area sites sampled for this work, with insert showing location of the Napa and Sonoma County sites. C. *Aphrophora* permutata nymphs on soap plant (*Chlorogalum pomeridianum*) at a sampled vineyard site in Napa County. D. *A. permutata* adult, Dry Creek Road, Napa County, 200m, with permission of Kim Wagner, CC BY-NC 4.0.

### An enigmatic vector of Pierce’s Disease

Previous research to identify potential vectors of Pierce’s Disease in Napa and Sonoma counties (California, USA) has uncovered substantial populations of a spittlebug of the genus *Aphrophora* capable of transmitting *X. fastidiosa* to grapevines (Beal et al., 2021). Initial morphological identification indicates this vector is Aphrophora permutata Uhler (Fig. 1C–D), consistent with the identification given by DeLong & Severin (1950) in their early studies of vectors of Pierce’s Disease (please see Supplemental Note S1 for our discussion on the taxonomic identities of these insects).

Kelson (1964) described the life cycle of A. permutata from Tilden Regional Park in the hills of east Berkeley, a region that comprises steep ridges and creek-drained valleys supporting oak woodland, coastal scrub, chaparral, grassland, and riparian forest. Nymphs feed on understory perennial herbs and vines. The adults move to overstory pine hosts (*Pinus* spp.) where they lay eggs in the needles. Each spring the nymphs hatch from pines, fall into the understory herbs, and repeat the life cycle. By contrast, the vineyards of Napa and Sonoma counties are intensively managed monocultures on comparatively flat terrain, with a simplified understory and regular insecticide inputs. Therefore, the populations of *A. permutata* that live in the vineyards in Napa and Sonoma County do not have immediate access to pine trees (Beal et al., 2021), which seems crucial for their lifecycle. This raises the question – are these vineyard spittlebugs *A. permutata* with a drastically different life cycle or, alternatively, a morphologically indistinguishable yet genetically distinct cryptic species?

In this study we use whole genome sequencing to unravel the taxonomic identity of the vineyard spittlebugs. Specifically, we: 1) genetically compare vineyard *Aphrophora* sp. in relation to *A. permutata*; 2) resolve the population structure of this enigmatic spittlebug across six vineyards in Napa and Sonoma, and investigate possible drivers of that structure including plant host usage, geography, and endosymbiont infection; and 3) determine how these Aphrophora have responded to selection pressures of insecticide usage in the vineyards.

## Materials and Methods

### Specimen collection

This study encompasses 48 samples from various sources over the past decade. Through the assistance of University of California Cooperative Extension personnel, samples were collected from three commercial vineyards in Sonoma County and three in Napa County (Fig. 1A–B; Supplemental Table S1). We also sampled three quasi-natural locales (Fig. 1A–B). The first (PM), along the Pine Mountain Fire Road in the Marin County highlands, consisted of Douglas fir (*Pseudotsuga menziesii*) and Monterey pine (*Pinus radiata*) scattered among grass-herb meadows. The second (CD) was composed of roadside Douglas fir trees along a rural section of Calistoga Road in Sonoma County. The third (TI) consisted of Monterey pine and Douglas fir trees in forested areas in Tilden Park in the Berkeley-Oakland Hills, the site of Kelson’s (1964) original studies of the *A. permutata* life cycle. Details, including host plants where known, life stage, and storage appear in Supplemental Table S1.

### DNA sequencing, quality control, and Population inference using Autosomal genomes

*We note that, for all the following computational analyses, additional flags and options outside of defaults and not included in the main methods text can be found in Supplemental Table S2*.

DNA extraction was completed using one leg from adults and two legs from nymphs with the ZymoBIOMICS DNA Miniprep Kit according to the manufacturer’s protocol. Library preparation and 150 bp genomic Illumina sequencing was conducted by Azenta Biotech on a NovaSeq X□+. We generated 10 GB of data for each sample to obtain ∼1-2x coverage per sample. Using BWA’s default implementation, we mapped the reads, as previously trimmed with Trimmomatic v0.36 (Bolger et al., 2014) by the manufacturer, to the reference genome of *A. alni* (Griffiths et al., 2024). We marked and removed duplicates with Picard (Broad Institute, 2023). We filtered out sex chromosomes from population structure analyses, for which many of our sampled males would have <1X coverage, and calculated genotype likelihoods for each individual only on autosomal chromosomes (nuclear genomes) using ANGSD v.0.921 (Korneliussen et al., 2014). Linked sites were identified and pruned with ngsLD v1.2.1 (Fox et al., 2019).

We conducted a Principal Components Analysis (PCA), with PCangsd (Meisner & Albrechtsen, 2018), using the unlinked sites as input, and visualized the centroids for three visually distinct genetic clusters. We also used the unlinked sites to generate admixture models with NGSadmix (Meisner & Albrechtsen, 2018), assuming 1-8 ancestral populations (K). For each level of K, we conducted 10 independent runs and assessed the optimal K by comparing the mean log-likelihood across replicates, the Evanno ΔK statistic (Evanno et al., 2005), and Akaike Information Criterion (AIC) based on the best log-likelihood-replicate likelihood for each level.

To further quantify the genetic differentiation in our samples we calculated the pairwise global F_ST_ between each of the three major sampling localities (Napa, Berkeley, and Sonoma) and between the three predicted genetic clusters identified in the autosomal PCA. Note that PM and CD samples were not included here because the number of samples is less than 5. We estimated the folded 2D Site Frequency Spectrum (SFS) across autosomes for each population pair with `realSFS -fold 1` from ANGSD (Korneliussen et al., 2014) as the F_ST_ prior. We then computed the weighted and unweighted F_ST_ estimates with `realSFS fst index` and `realSFS fst stats2`. To control for possible bias introduced by variable coverage among our samples, we tested whether coverage varied among clusters or correlated with PC scores, ancestry proportions or ancestry entropy, and repeated the PCA and admixture analyses without the two lowest-coverage individuals (see Supplemental Methods 1).

### Population inference using Mitochondrial genome

We modeled population structure from the mitochondrial genome separately from the nuclear genome. We called variants on the mitochondrial genome with bcftools v1.19 (Danecek et al., 2021; Li, 2011), setting minimum quality and map quality to 20, and retaining both variant and invariant sites. We then generated per-individual consensus sequences with bcftools `consensus`, masking positions absent from the VCF and calling missing genotypes as N, and realigned the sequences with MAFFT v7 (Katoh & Standley, 2013).

We computed genetic distances between individuals as the raw proportion of differing sites, with APE v5.8.1 (Paradis et al., 2004) and then conducted a principal coordinates analysis (PCoA) on the distance matrix. To allow visual comparison between the nuclear and mitochondrial genomes, a centroid was plotted for each of the three autosomal genetic clusters to assess whether mitochondrial structure recovers the autosomal grouping. We extracted biallelic SNPs from the mitochondrial genome and conducted a clustering analysis with *sparse non-negative matrix factorisation* (*smnf*; Frichot et al., 2014) from LEA v 3.16.0 (Frichot & François, 2015). We again tested K 1-8, with 10 replicates per K. We selected the best model with *smnf*’s cross-entropy criterion, an Evanno-style ΔK computed on cross-entropy, and the spread of cross-entropy across replicates.

To quantify the variation in mitochondrial genomes, we used the Φ_ST_ statistic. We computed Φ_ST_ from the sums of squared pairwise distances following Excoffier et al. (1992) globally across autosomal clusters, and pairwise between the three population comparisons. Significance was assessed by conducting 1000 permutations of cluster labels and recomputing Φ_ST_ for each permutation, with a Bonferroni correction applied to all three pairwise p-values.

### Comparisons within the Genus Aphrophora using Cytochrome oxidase 1 (COI)

To understand the overall relationship of the vineyard *Aphrophora* to congeners, we evaluated the relationships using the gene fragment COI, extracted from the mitochondrial genomes. We downloaded all available COI sequences from NCBI GenBank for *Aphrophora* which included a species identification (Supplemental Table S3). Sequences were aligned using MUSCLE (Edgar, 2004, 2022) and a phylogenetic tree was recovered using IQTREE3 (Wong et al., 2025). We chose *Philaenus spumarius* as an outgroup for tree reconstruction. Distantly related species *Aphrophora alni*, *A. quadrinotata*, *A. pectoralis*, *A. salicinia*, *A. canadensis*, *A. corticea*, *A. sartogensis*, *A. parallela* and *A. cribrata*, as identified by the tree, were not used in haplotype network analysis. The haplotype network was generated in PopART (Leigh & Bryant, 2015) using the minimum spanning network (Bandelt et al., 1999).

### Inference of Demographic History

We used GADMA2 (Noskova et al., 2020, 2023) to infer the demographic history between the two cryptic populations of *A. permutata* as identified by admixture analyses. As Clusters 1 and 3 are of comparable sizes, and Clusters 2 and 3 belong to the same genetic population, we included only individuals that localized to Cluster 3 as representatives from our second A. permutata population. We also excluded individuals with coverage below ∼1x from both populations. Because selection (Johri et al., 2021) and repetitive elements (Patil & Vijay, 2021) can heavily bias demographic inference, we estimated the folded SFS (with *A. alni* as the ancestor) for both of these populations with ANGSD, excluding repetitive genomic elements, and coding sequences (with a 10 Kb buffer around coding sequences). We only included sites from autosomes, requiring at least 80% of the individuals per site and projected the SFS to haploid genomes by hypergeometric projection (Marth et al., 2004; Gutenkunst et al., 2009).

Using the moments engine (Jouganous et al., 2017) we fit three competing models to the joint SFS. The first contained only a split between the two populations; the second and third added continuous asymmetric and symmetric gene flow respectively. For each model, we conducted 50 independent replicates, each with a different random initialisation and concluding with a local search (`BFGS_log`) from the best solution reached by the genetic algorithm. The replicate with the highest composite log-likelihood is taken as that model’s best fit (θ).

To bootstrap the model, we divided the autosomes into 10 Mb windows, a conservative choice exceeding the expected scale of linkage, and retained windows only if they contained ≥ 10Kb of putatively neutral base pairs (non-repetitive, non-genic, and not gene-adjacent) giving 157 blocks from 1,161 windows and > 500 million bp of sequence in total. We computed the joint SFS once per block and created 100 bootstrap replicates formed as the sum of blocks resampled with replacement. These bootstrap replicates were provided to GADMA2 to estimate the Composite Likelihood Akaike Information Criterion (CLAIC; Coffman et al. 2016; Noskova et al. 2020) for each model’s best replicate and compare the model fits, and independently calculated AIC for each model.

We then computed confidence intervals (CI) for the model parameters on the CLAIC-selected best demographic model. We re-ran local optimisation from the inferred optimum θ[ on each of the 100 bootstrap replicates. To scale the model parameters into years, we assumed a genome-wide mutation rate of 2.9 × 10_J_J and a generation time of one year (*A. permutata* is univoltine [Kelson 1964]).

### Testing for drivers of population structure

We tested how factors like host plant usage, insecticide resistance, and endosymbiont infection could drive population structure by looking for the outlier loci in the genome and their function.

### Identification of outlier loci

To identify loci under selection in each autosomal cluster, we conducted genome-wide scans using non-overlapping 50kb windows, and calculated F_ST_ between each pair of autosomal clusters recovered from PCA analysis. We computed the windowed pairwise F_ST_ estimates using `realSFS fst stats2` from ANGSD (Korneliussen et al., 2014) using the same 2D SFS priors as the global analysis. We then computed Population Branch Excess (PBE; Yassin et al., 2016) for each of the three autosomal clusters from ∼28,000 non-overlapping F_ST_ estimates. We considered windows in the upper 1% of the PBE distribution per cluster as candidates for selective sweeps, per Shpak et al. (2025). We also considered windows in the bottom 1% of all three pairwise F_ST_ distributions, and in the bottom 1% of F_ST_ windows between Clusters 1 and 3 (which both predominantly inhabit vineyards) as windows containing putatively adaptive shared loci due to uniform selection, although we acknowledge such tests do not definitively identify selection.

### Testing for host plant usage and insecticide resistance

To determine the functions of genes in outlier regions, we functionally annotated the previous annotation of *A. alni* (Griffiths et al., 2024) against eggNOG 5.0 (Huerta-Cepas et al., 2019) with eggNOG-mapper v2.1.12 (Cantalapiedra et al., 2021) using the DIAMOND protein aligner (Buchfink et al., 2015) and restricting orthology to Insecta. We considered two categories of genes in our outlier regions. First, we identified chemosensory (Supplemental Table S4), including olfactory gene families, as they can be highly tied to host plant usage (Matsuo et al., 2007; McBride, 2007; C. Smadja & Butlin, 2009; C. M. Smadja et al., 2012), and to the evolution of prezygotic mating barriers (C. Smadja & Butlin, 2009).

Second, insecticide usage can exert intense selection pressures, resulting in selective sweeps and rapid evolution (P. J. Daborn et al., 2002; Hawkins et al., 2019; Karasov et al., 2010; Labbé et al., 2007). We therefore reasoned that adaptation to insecticides could drive population structure in our sampled *A. permutata*, and that populations could have unique resistance signatures. To assess this we identified known insecticide-resistance genes (Supplemental Table S4) in our windows of interest.

### Endosymbiont infection

As the presence of the bacterial endosymbiont *Wolbachia* is hypothesized to create reproductive barriers between infected and uninfected populations in spittlebugs through cytoplasmic incompatibility (Lis et al., 2015; Seabra et al., 2021), we screened our samples for bacterial endosymbionts. For all individuals we considered candidate endosymbiont reads where at least one mate was unmapped, or where mapping quality to the reference genome was < 10 (samtools view -e; Danecek et al., 2021). To identify putative symbionts, we created a reference panel of eight host endosymbiont genomes (Supplemental Table S5). We included the reference *A. alni* genome in the screen, to differentiate reads which may preferably map to *Wolbachia* genome insertions in the host genome (Dunning Hotopp et al., 2007). We mapped reads against our reference panel with BWA-MEM (Li & Durbin, 2009), retaining paired alignments with a mapping quality score of at least 30, and used samtools depth (Danecek et al., 2021) to calculate coverage across the reference panel.

We calculated coverage breadth as the fraction of the endosymbiont genome coverage of at least 1X, and summarized coverage evenness as the Gini coefficient of mean read depth across non-overlapping 1kb windows spanning each endosymbiont reference genome (Gini & Pearson, 1912; Lorenz, 1905). We considered each individual to be infected with an endosymbiont at coverage breadth > 0.30 and G < 0.60. To control for possible bias introduced by sequencing depth, we compared the mean depth of coverage on the host genome between individuals putatively infected with an endosymbiont to uninfected individuals with the Wilcoxon rank-sum test.

We further tested whether endosymbiont infection (Wolbachia and Rickettsia) was associated with sampling locality or climate, using site-level Fisher’s exact tests, Mann–Whitney U tests, and rank correlations, together with permutation tests of spatial structure (20,000 iterations each). Full details, including the thermal indices (Supplemental Table S6) derived from Open-Meteo climate data (1995–2024) and the permutation procedures, are given in Supplemental Methods 2.

## Results

### Population Structure Analysis

Our autosomal PCA uncovered differentiation between the Berkeley population and all other sampled populations across PC2, with no evidence of structure between the geographic Napa and Sonoma (Fig. 2A) populations. We observed three distinct clusters in the PCA, Cluster 1 (PC1 > 0.1) was comprised of individuals from Berkeley, Napa and Sonoma, Cluster 2 (PC1 < −0.05; PC2 > 0.1) contained only individuals from Berkeley, and Cluster 3 (PC1 < 0; PC2 < 0) was made up of individuals from Napa, Sonoma (including CD) and PM (Fig. 2A and Supplemental Fig. S1). Coverage did not differ among clusters and was uncorrelated with PC and ancestry component (Supplemental Table S7). Population structure did not change when the two PM individuals (which had the lowest coverage at <0.7x; see Supplemental Table S1 for per-sample coverage) were dropped from the analysis (Supplemental Figs. S2–S3).

**Fig. 2.**
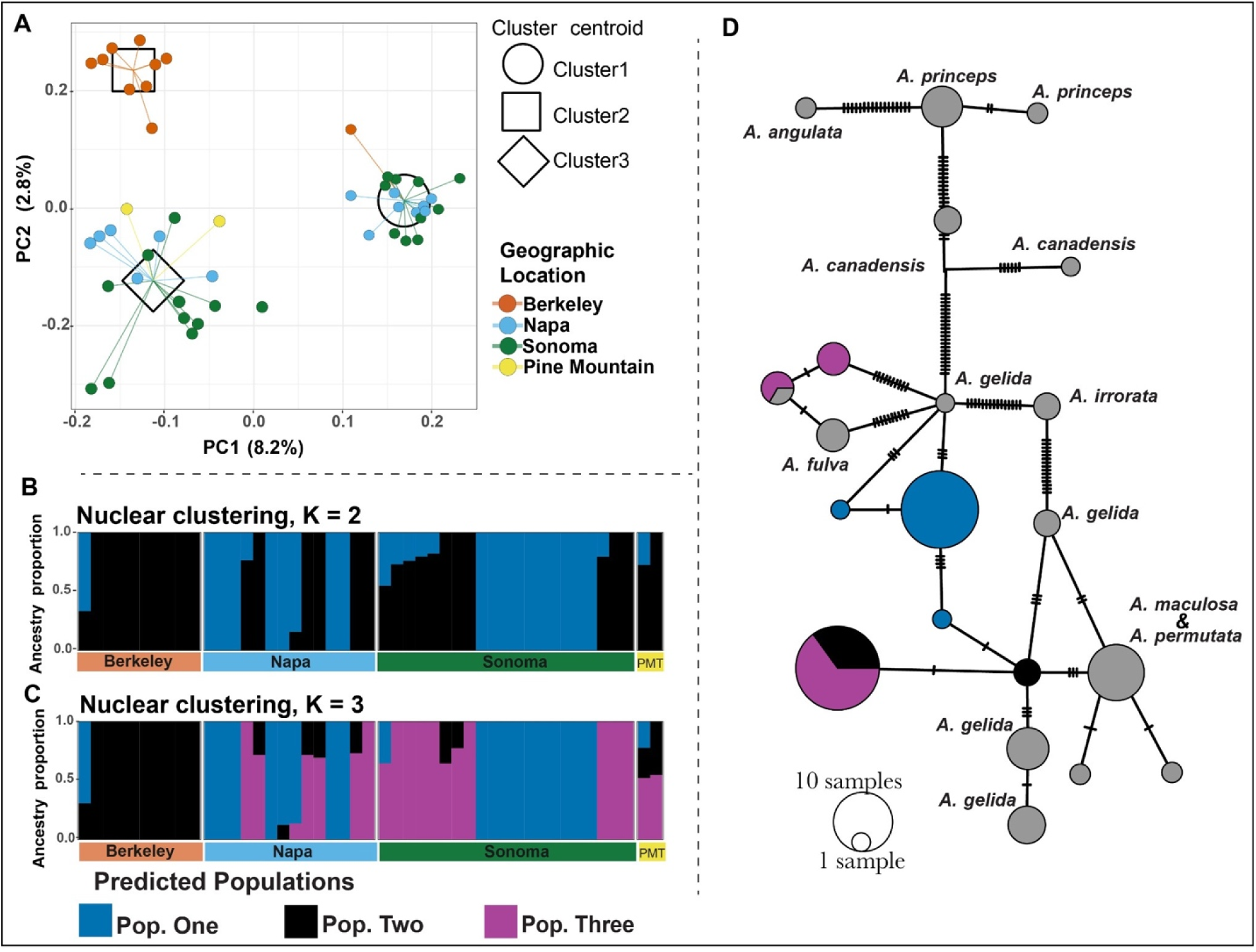
Autosomal analysis. A) Principal Components Analysis (PCA) from autosomal SNPs. B) Admixture components from autosomal SNPs at the best-fitting K=2 admixture model. C) Admixture components from autosomal SNPs at K=3 model admixture. Individuals are coloured according to sampling locality (Berkeley, Napa, Sonoma and Pine Mountain), with the centroids for individuals assigned to Clusters 1, 2 and 3 in the autosomal PCA demarcated by a circle, square and diamond respectively. D) Autosomal genome clusters mapped onto mitochondrial COI haplotype network of A. permutata and closely related species.

Both Evanno’s ΔK and AIC supported K=2 admixture components for autosomal nucleotide likelihoods (Supplemental Fig. S4A–C). At K=2 autosomal components were approximately evenly divided in both Napa and Sonoma, while Berkeley was strongly skewed towards autosomal component 1 (Fig. 2B). At K=3, autosomal component 2 was primarily found in Berkeley, while the other two sampling localities contained autosomal components 1 and 3 at approximately equivalent levels (Fig. 2C). We identified little structure between geographically sampled populations (FST = 0.04 between Berkeley and both Napa and Sonoma; F_ST_ = 0.01 between Napa and Sonoma; Table 1). Of the three autosomal genetic clusters, Clusters 2 and 3 had the least population structure (F_ST_ = 0.05), Clusters 1 and 2 having the strongest structure (F_ST_ = 0.08), and Clusters 1 and 3 also showing moderate structure (F_ST_ = 0.06; Table 1).

**Table 1.**
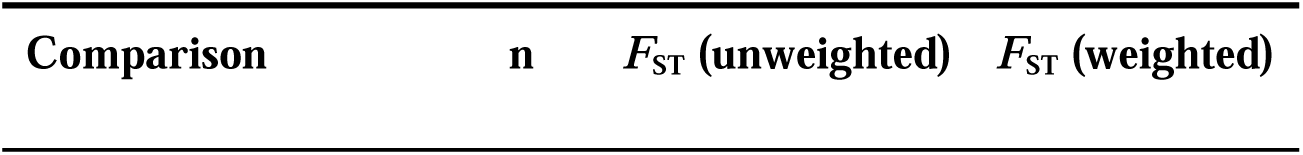
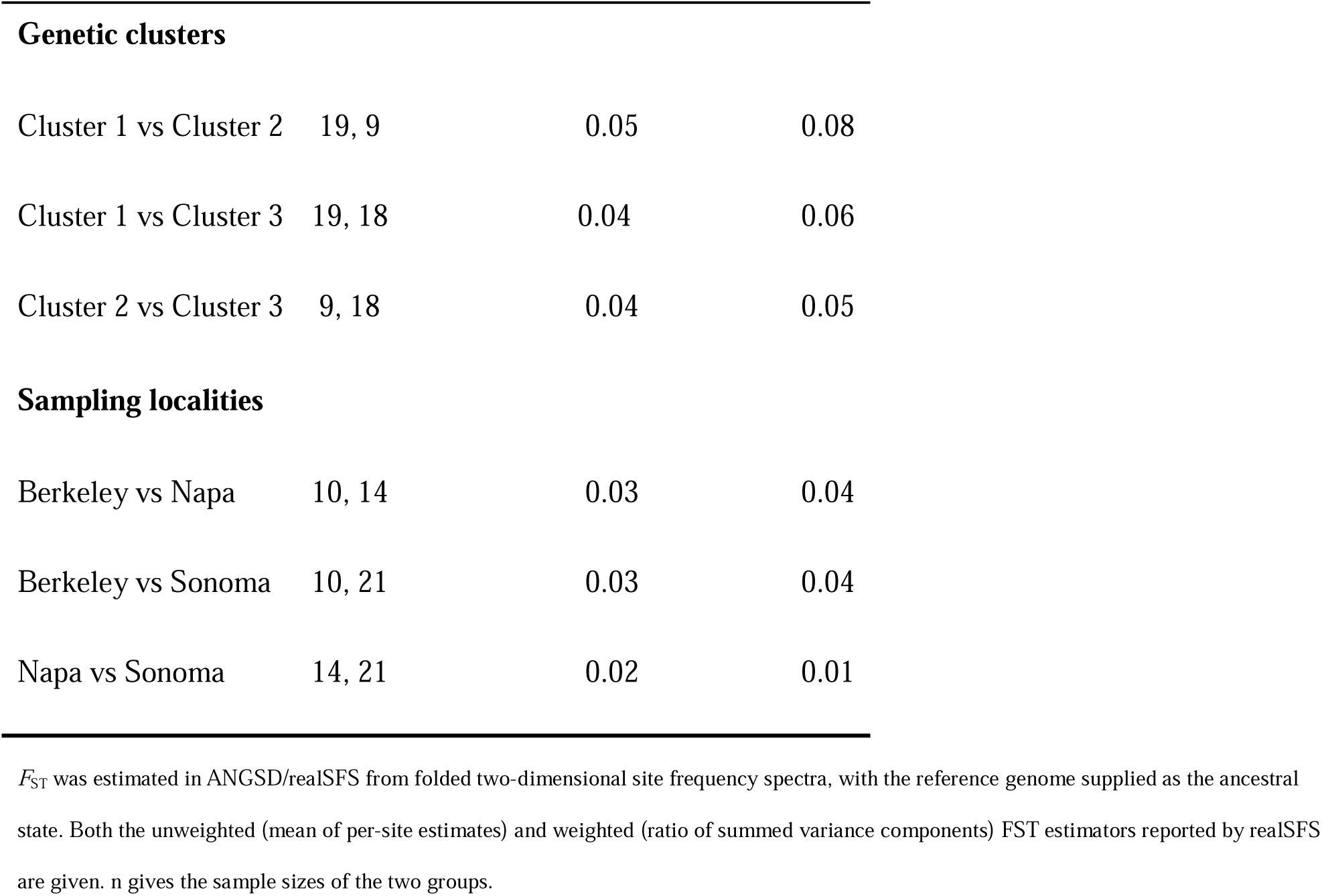
Autosomal genetic differentiation (*F*_ST_) between genetic clusters and between populations of Aphrophora.

| Comparison | n | $F_{ST}$ (unweighted) | $F_{ST}$ (weighted) |
| --- | --- | --- | --- |
| **Genetic clusters** |  |  |  |
| Cluster 1 vs Cluster 2 | 19, 9 | 0.05 | 0.08 |
| Cluster 1 vs Cluster 3 | 19, 18 | 0.04 | 0.06 |
| Cluster 2 vs Cluster 3 | 9, 18 | 0.04 | 0.05 |
| **Sampling localities** |  |  |  |
| Berkeley vs Napa | 10, 14 | 0.03 | 0.04 |
| Berkeley vs Sonoma | 10, 21 | 0.03 | 0.04 |
| Napa vs Sonoma | 14, 21 | 0.02 | 0.01 |

$F_{ST}$ was estimated in ANGSD/realSFS from folded two-dimensional site frequency spectra, with the reference genome supplied as the ancestral state. Both the unweighted (mean of per-site estimates) and weighted (ratio of summed variance components) $F_{ST}$ estimators reported by realSFS are given. n gives the sample sizes of the two groups.

All but one individual assigned to Cluster 1 in the autosomal PCA clustered closely together in the mitochondrial PCoA, distal from the other samples by PCo1, whereas those from Cluster 2 and Cluster 3 in the autosomal PCA were split across two clusters on PCo2 (Fig. 3A). A third cluster, distal across PCo2 only contained individuals from Sonoma (Fig. 3A). Our Evanno-style ΔK marginally supported K=2 over K=3 mitochondrial clusters, although K=3 was preferred by mean cross-entropy, and only moderately underperformed K=4 in replicate spread (Supplemental Fig. S5A–C). At K=2 and K=3 all Berkeley and Pine Mountain replicates were most likely to have inherited mitochondrial component two (Fig. 3 B-C). Napa and Sonoma were split between components one and two at K=2, and Sonoma was the only population to contain mixture three at K=3 (Fig. 3B-C). Individuals falling within autosomal Cluster 1 are nearly completely differentiated from other autosomal clusters in the mitochondrial genome (ΦST = 0.958 and 0.833, p < 0.001; Table 2). Autosomal Clusters 2 and 3 are non-significantly differentiated from each other (ΦST = 0.167, p = 0.09; Table 2).

**Fig. 3.**
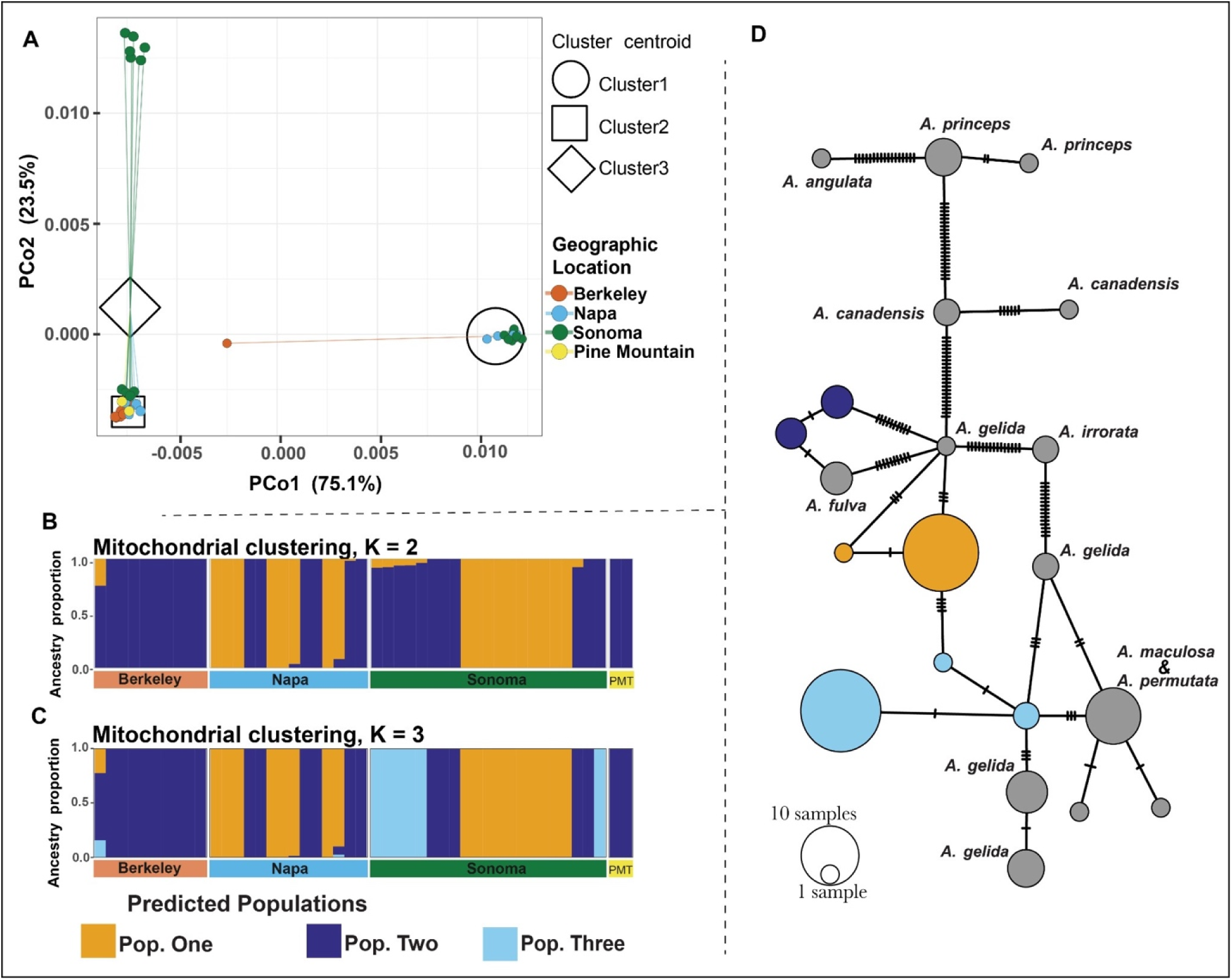
Mitochondrial genome analysis. A) Principal Coordinates Analysis (PCoA) from mitochondrial SNPs. B) Mitochondrial clustering identified with sparse non-negative matrix factorisation (smnf) in the best fitting K=2 model. C) Mitochondrial clustering identified with sparse non-negative matrix factorisation (smnf) in the K=3 model. D) Mitochondrial genome clusters mapped onto mitochondrial COI haplotype network of A. permutata and closely related species.

**Table 2.** Mitochondrial differentiation (**Φ**_ST_) among the three autosomal genetic clusters of Aphrophora.

| Comparison (autosomal clusters) | n | $\Phi_{ST}$ | P | P (Bonferroni) |
| --- | --- | --- | --- | --- |
| Global (3 clusters) | 46 | 0.828 | < 0.001 | — |
| Cluster 1 vs Cluster 2 | 19, 9 | 0.958 | < 0.001 | < 0.003 |
| Cluster 1 vs Cluster 3 | 19, 18 | 0.833 | < 0.001 | < 0.003 |
| Cluster 2 vs Cluster 3 | 9, 18 | 0.167 | 0.029 | 0.087 |

Analysis of molecular variance on 13,204 bp of mitochondrial sequence from 47 individuals, using uncorrected p-distances with pairwise deletion of missing data. Significance was assessed by 1,000 permutations of cluster labels; the smallest attainable P-value is 1/1001 and is reported as < 0.001. Pairwise P-values were Bonferroni-corrected for three comparisons; the global test is a single omnibus test and is reported uncorrected.

Our sampled COI sequences fell within a clade which included *A. fulva, A. gelida*, *A. maculosa*, and *A. permutata,* with none of the species in this clade recovered as monophyletic (Supplemental Fig. S6). Autosomal Cluster 1 largely shared a COI haplotype, while the haplotypes for the other two populations were more diffuse (Fig. 2D). COI haplotypes closely followed mitochondrial population structure (Fig. 3D) recovered using PCoA.

### Demographic modeling

CLAIC could not be computed for our no post-split migration model, because its Godambe information matrix was singular at every finite-difference step size attempted; it was nonetheless rejected, as it was 2,022 log-likelihood units worse than the other models, and was heavily penalized by AIC (Supplemental Table S8). The symmetrical and asymmetrical migration models were statistically indistinguishable, differing by only 0.05 log-likelihood and 0.02 CLAIC units. We considered the symmetrical migration model as the preferred model as the most parsimonious choice (*k* = 7 vs. 8), a choice supported by its lower AIC (2,528.91 vs. 2,530.79; Supplemental Table S8).

The demographic model (Fig. 4A) generally fits the empirical data (Fig. 4B). Within this model, the ancestral effective population size (N_e_) was 285,300 (95% CI: 191,700 – 319,500), lasting nearly 100 thousand years (95% CI: 76,700 – 114,300; Table 3). The two populations split nearly evenly (ancestral population founding Cluster 3=50.8%; 95% CI: 41.1-59.3%) between 39,050 and 87,050 years ago (point estimate 44,970; Table 3). While Cluster 3 (N_e_ = 476,000) is estimated to currently have a smaller population size than Cluster 1 (Ne = 676,800), these estimates have overlapping confidence intervals and are not significantly different. Gene flow between the two lineages was substantial and equivalent in rate, amounting to 2.1 (95% CI: 1.8–2.9) effective migrants per generation into Cluster 3 and 3.0 (95% CI: 2.4–3.8) into Cluster 1 (Table 3).

**Fig. 4.**
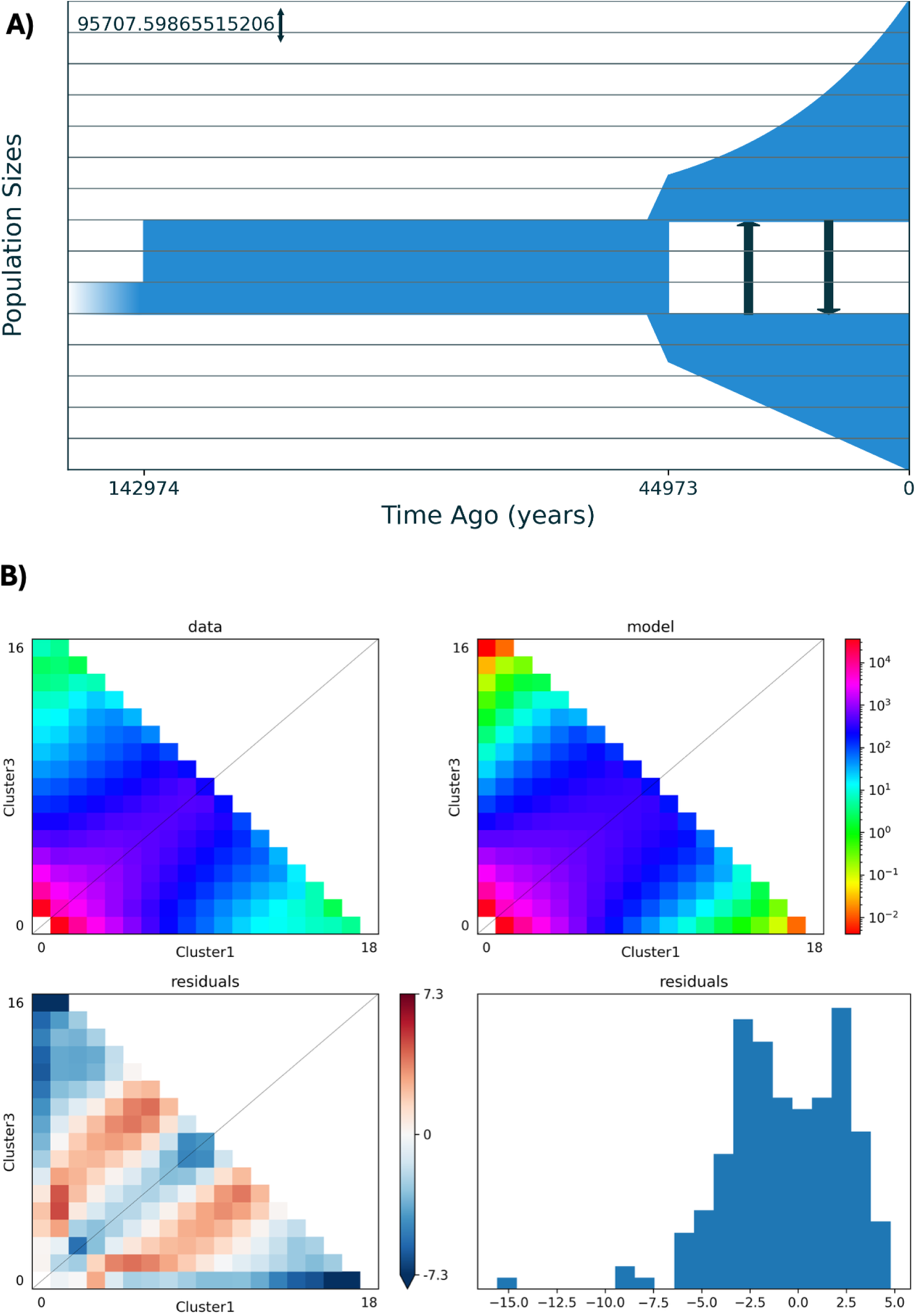
Demographic history of sampled A. permutata. A) Demographic history, with symmetrical gene flow represented with black arrows. The root ancestral effective population size (NA _≈_ 95,700), the reference against which all sizes in Table 3 are scaled, is shown with double-sided arrows for scale. B) Model fit compared to the observed site frequency spectrum.

**Table 3.** Demographic history of the Cluster 3–Cluster 1 divergence under the selected model (split with symmetric gene flow), in biological units. Values are the maximum-composite-likelihood estimate with 95% percentile confidence intervals from 100 block-bootstrap replicates. Parameters correspond to the epochs shown in Fig. 4A.

|  | Estimate | 95% CI |
| --- | --- | --- |
| *Before divergence* |  |  |
| Ancestral effective population size (ancestral epoch, after expansion from NA) | 285,300 | 191,700 – 319,500 |
| Duration of the ancestral epoch (years) | 98,000 | 76,700 – 114,300 |
| *Divergence* |  |  |
| Time since divergence (years ago) | 44,970 | 39,050 – 87,050 |
| Ancestral population founding Cluster3 (%) | 50.8 | 41.1 – 59.3 |
| *Since divergence* |  |  |
| Cluster3 effective population size | 476,600 | 372,100 – 544,700 |
| Cluster 1 effective population size | 676,800 | 469,400 – 757,400 |
| Migrants per generation into Cluster3 | 2.1 | 1.8 – 2.9 |
| Migrants per generation into Cluster 1 | 3.0 | 2.4 – 3.8 |

Gene flow is symmetric in rate. The number of migrants differs between the two directions only because the receiving populations differ in size.
Intervals were computed by recalculating each quantity within every bootstrap replicate, using that replicate's own $\theta$ , so uncertainty in the scaling is carried through. They exclude uncertainty in the mutation rate and generation time, both treated as fixed, and uncertainty about the model itself.

### Drivers of population structure

#### Chemosensory and Insecticide Resistance Gene Families

Four chemosensory genes fell within PBE top-1% windows: an unnamed ionotropic receptor in Cluster 1, as well as an SNMP/CD36 protein, a chemosensory protein (*Csp*) and a dual CSP/OBP pheromone-binding gene of the A10/OS-D family in Cluster 2. One gustatory receptor fell in an undifferentiated genomic region (bottom 1% F_ST_) between autosomal Clusters 1 and 3.

Each autosomal genetic cluster contained candidate insecticide-resistance genes within regions showing signatures of recent positive selection (also referred to as selective sweeps; Supplemental Table S9). These fell into two functional groups: insecticide target sites (nicotinic acetylcholine receptors) and detoxification enzymes (Cytochrome P450 monooxygenases, carboxylesterases, ABC transporters, and UDP-glycosyltransferases). Three nicotinic acetylcholine receptors fell within swept regions in Cluster 2, and two others in Cluster 1. Cluster 3 contained four Cytochrome P450 monooxygenases, while a single carboxylesterase occurred in both Clusters 1 and 3. One ATP-binding cassette transporter was recovered in Cluster 2 and three in Cluster 3, and Clusters 2 and 3 each contained one UDP-glycosyltransferase (Supplemental Table S9).

Three Carboxyl/cholinesterase genes were found in windows of low structure (bottom 1% F_ST_ windows) across all three genetic clusters, which may imply uniform selection (Supplemental Table S10; the two loci on chromosome 9 are adjacent paralogues). Five candidate insecticide-resistance genes were found in regions of low structure between Clusters 1 and 3, including two nicotinic acetylcholine receptors, two ATP-binding cassette transporters, and one UDP-glycosyltransferase.

#### Endosymbiont infection

Only the wMel strain of *Wolbachia* supergroup A (n=13) and *Rickettsia* (n=8) showed evidence of active infection in our sampled individuals (Supplemental Table S11; Fig. 5; Supplemental Fig. S7). Sequencing coverage of the host did not affect detection of *Wolbachia* (mean infected = 1.40x; mean uninfected = 1.49x; W = 211; p = 0.99) or *Rickettsia* (mean infected = 1.41x; mean uninfected = 1.48x; W = 147; p = 0.81). Notably, both PM individuals, which had the lowest average host coverage (< 0.70) were positive for *Wolbachia* and one of these individuals was also infected with *Rickettsia*, suggesting we were not missing infection due to low coverage. *Wolbachia* infection was strongly structured among sites, occurring in all individuals from Berkeley and PM and one from Napa (permutation across individuals, variance in site frequency = 0.167, *p* < 0.001; reported descriptively, as this individual-level test is pseudoreplicated). *Rickettsia* showed no such structure, occurring as scattered low-frequency infections at four sites spanning both habitat types (variance = 0.040, *p* = 0.168).

**Fig. 5.**
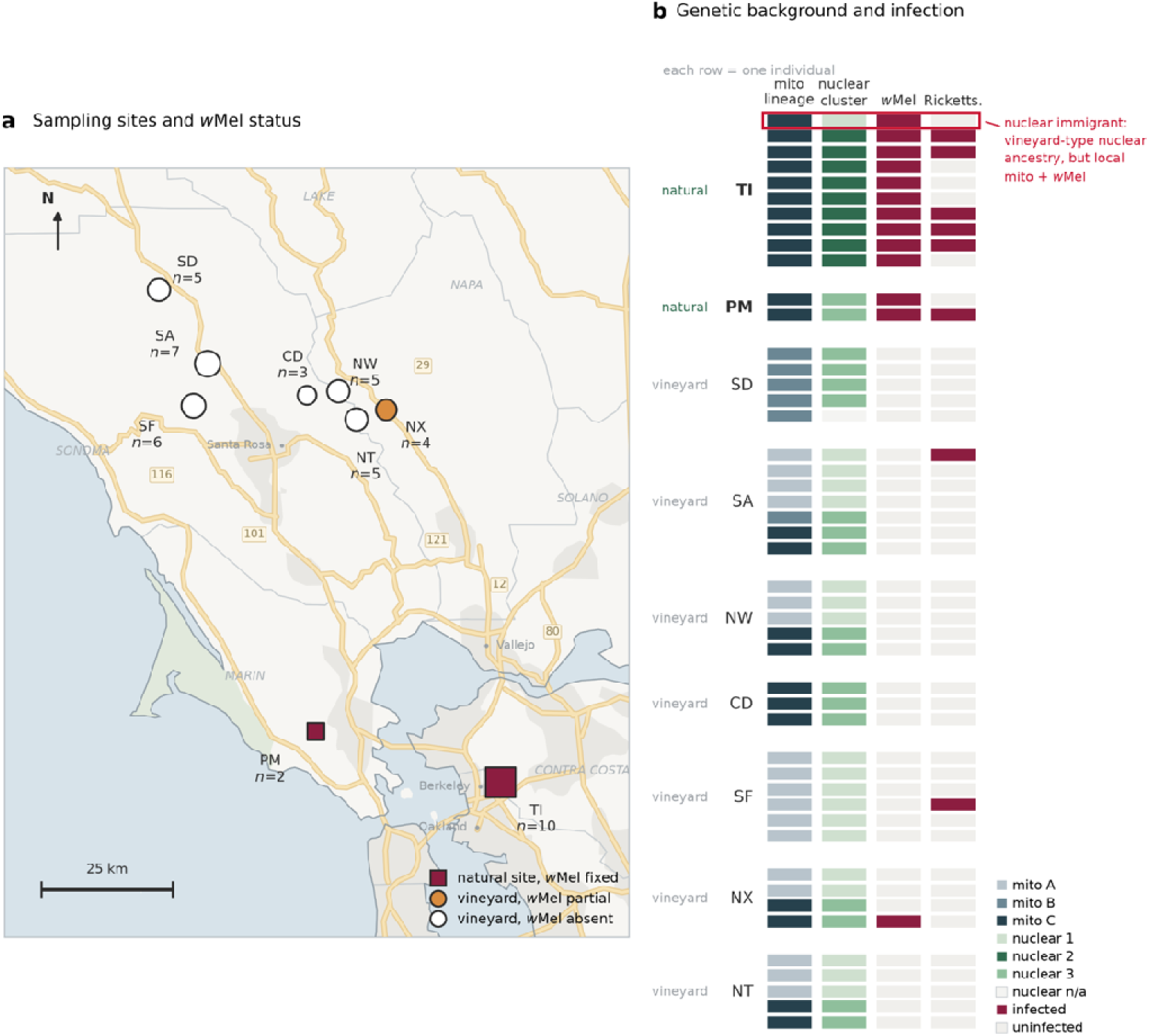
Endosymbiont infection: Wolbachia is structured by site, not by geography. (**a**) Nine sampling localities in the North Bay / East Bay, California (n = 47, low-coverage whole-genome sequencing). Squares mark the three quasi-natural sites, circles the six vineyards (two-letter codes); fill gives site-level wMel status (fixed, partial, absent) and marker area sample size, printed beside each site. Coordinates are approximate (∼200 m). (**b**) One row per individual, grouped by site: mitochondrial lineage, autosomal cluster, and wMel and Rickettsia infection. The outlined individual carries vineyard-type autosomal ancestry with the local mitochondrion and wMel.

*Wolbachia-*infected individuals were assigned mostly to autosomal Cluster 2, but Clusters 1 and 3 also contained infected individuals owing to the universal infection in Berkeley (Fig. 5; Supplemental Fig. S7). All infected *A. permutata* had mitochondrial Cluster 3 ancestry, although this lineage was not universally infected (Fisher’s exact test, p < 0.01). Mitochondrial background is therefore necessary but not sufficient to explain the distribution: within Cluster 3 alone, infection remained strongly structured by site (permutation, variance = 0.194, *p* < 0.01), with uninfected Cluster-3 individuals occurring at vineyard sites. The association is thus with site of origin rather than with lineage membership. *Rickettsia* was not associated with mitochondrial background (OR = 3.88, *p* = 0.137).

Neither spatial nor environmental variables predicted infection. Infected localities showed no spatial clustering (mean pairwise separation 57.2 vs. 44.5 km for randomized triplets; permutation p = 0.686), and elevation had no site-level effect (Mann–Whitney U = 8.0, p = 1.00), with the two highest sites entirely uninfected. No thermal metric distinguished infected from uninfected sites; mean May–October temperature, for example, was uncorrelated with infection frequency (U = 6.0, p = 0.548; Spearman ρ = −0.418, p = 0.263). The only infected vineyard was also the warmest site, contradicting the hypothesis that thermal stress limits endosymbiont presence—though excluding this outlier would render the association significant (ρ = −0.756, p = 0.030). The ten thermal metrics yielded nearly identical rankings, precluding their use as competing hypotheses.

The lone Berkeley individual with autosomal Cluster-1 ancestry still carried the local mitochondrial lineage and wMel infection. Site-level infection appeared elevated at the two quasi-natural sites relative to vineyards, but not significantly (Fisher’s exact test, p = 0.083), and habitat type is likely confounded with latitude. Two further indicators of restricted dispersal—lower mitochondrial lineage richness and reduced within-site mitochondrial dispersion at non-agricultural sites—were not supported (U = 3.0, p = 0.252 and p = 0.333, respectively).

## Discussion

We had three aims here: to clarify the relationship between *the vineyard Aphrophora sp. and A. permutata;* to resolve the population structure of this spittlebug across six vineyards in Napa and Sonoma counties and identify its drivers; and, lastly, to determine how these insects have responded to insecticide use across our sampled localities. Our results clearly show that the vineyard *Aphrophora* is in fact *A. permutata,* likely with a different life history. We find concrete evidence that the Napa and Sonoma vineyard populations are the same species as those found in quasi-natural habitats of Berkeley, Pine Mountain, and Calistoga Road (F_ST_ < 0.05), yet they comprise two distinct, largely sympatric populations of *A. permutata*. Below we consider why geography has not driven this population structure, alongside the possible roles of host plant usage, endosymbionts, and insecticide resistance in adaptation to a human modified landscape, and what our findings imply for taxonomy and species delimitations in *Aphrophora*.

### Population structure in A. permutata

#### Neither geography nor habitat usage explains population structure

The San Francisco Bay (Fig. 1A–B) is a documented barrier to gene flow in other Bay Area taxa like amphibians and reptiles (Rissler et al., 2006; Reilly et al., 2015), yet we do not see any signatures of this structuring in our sampled *A. permutata* (Fig. 2A).

Dispersal between Napa and Sonoma, and Berkeley may have genetically homogenized these populations. Although many spittlebugs spend their adult lives in or near one locality (Bodino et al., 2021; Lago, Morente, et al., 2021), some can travel over 1 km in hours (Lago, Garzo, et al., 2021; Casarin et al., 2023) and may also “hitchhike” along major roads (Bajocco et al., 2023). *A. permutata* could therefore traverse the bay by flight or by hitchhiking. Alternatively, the full Golden Gate barrier may have opened only within the past ten thousand years (Atwater et al., 1977), which is likely insufficient time for strong population structure to accumulate.

The Mayacamas mountains separating Napa and Sonoma likewise show no population structure in *A. permutata* (F_ST_ = 0.01). While the California Coast range is documented to drive east – west differentiation across animals (Calsbeek et al., 2003), the Mayacamas mountains remain underexplored in this context. Intriguingly, *X. fastidiosa* is strongly differentiated between the Napa and Sonoma valleys (Vanhove et al., 2020). Given its low (∼5%) transmission rate by Aphrophora to grapevines in the laboratory (Beal et al., 2021), this contrast—strong structure in the pathogen, little in the vector—could arise in several ways. *A. permutata* may move between valleys often enough to homogenize its own populations but transmit *X. fastidiosa* too rarely to homogenize the pathogen, whose structure may instead be driven by leafhopper vectors. Alternatively genetic homogenization between these populations may be maintained through stepping-stone gene flow, in which case *X. fastidiosa* need not spread across the host’s entire genetic neighborhood.

Habitat type also fails to explain structure. Berkeley individuals separate slightly from Napa and Sonoma along PC2 (Fig. 2A), but this axis captures under 3% of genetic variation, and admixture, FST (Table 1), and mitochondrial analyses all place them with vineyard individuals in the third genetic cluster (Supplemental Fig. S1). Critically, individuals of mixed population origin occur in both quasi-natural and vineyard sites, including one Berkeley individual with largely Cluster-1 ancestry (Fig. 2A–D, Fig. 3A–D). Hence quasi-natural and agricultural habitats do not harbour any genetically distinct populations.

#### Host Plant Usage

It is clear from our collecting efforts (Supplemental Table S1) that *A. permutata* adults live on both Monterey pine and Douglas fir. On Pine Mountain Trail, *A. permutata* adults were abundant on a Monterey pine tree and a patch of Douglas fir trees separated by only 50 meters. This, alongside extensive collections from both host plants (Supplemental Table S1) indicates that both species are attractive hosts. In the vineyard areas sampled in Napa and Sonoma counties, pine trees are rare or absent but Douglas firs are often present and, in some circumstances, abundant, suggesting that Douglas fir could serve as the summer host for at least some of the *A. permutata* adults that disappear from vineyards during the summer dry period. The three *A. permutata* specimens we collected in August from Douglas fir on Calistoga Road give credence to this hypothesis.

Using more than one host plant or habitat type is not unusual for a spittlebug (Thompson et al., 2023; Thompson, 2026; Bodino et al., 2020; Beal et al., 2021; Supplemental Table S1). However, our results do not mean that host plant usage has never driven population structure in *A. permutata*. The landscape of possible host plants has changed drastically since the split of these populations (Table 3; Fig. 4), and especially since the arrival of humans. For example, while Douglas fir is native to Napa, Sonoma and Marin counties (The Gymnosperm Database, 2026), it was probably introduced into the Berkeley-Oakland area by artificial planting (Ertter & Naumovich, 2013). The Monterey pine’s natural distribution is limited to small enclaves on the California coast south of San Francisco and two small islands off Baja California (The Gymnosperm Database, 2026). It is introduced and considered invasive in the Berkeley-Oakland area (Ertter & Naumovich, 2013). This suggests that the *A. permutata* populations studied were likely native in Napa, Sonoma and Marin Counties, but introduced to the Berkeley-Oakland Hills in historical times, or that they historically used a different adult host plant.

#### Bacterial endosymbionts

*Wolbachia* does not appear to be a major driver of population structure, but its own distribution may track temperature or habitat in our system. Some strains of *Wolbachia* supergroup A (the strain identified here) have shifted host preferences towards cooler temperatures in Drosophila (Hague et al., 2020). As the only sites impacted by the coastal advection fog, Berkeley and Pine Mountain are expected to be cooler, on average, than sampling sites in Napa and Sonoma counties (Fischer et al., 2009; Iacobellis & Cayan, 2013; Johnstone & Dawson, 2010). Climate data bear this out: both average 2.6 °C cooler over the May–October activity window than the next coolest site. Because Pine Mountain and Berkeley are the only sampled localities well outside of agricultural reaches, we cannot discount the possibility that *Wolbachia* is selected against in a more heavily altered habitat. Because the two fog-influenced sites are simultaneously the coolest, the only non-agricultural, and the two southernmost localities, temperature, habitat and latitude cannot be separated in this design, but the high infection rates at Berkeley merit further consideration. As the identification of one *Wolbachia* infected individual at our warmest locality biases our modelling efforts, further sampling is needed to more robustly understand the relationship between *Wolbachia* infection and temperature.

Like *Wolbachia*, the maternally inherited endosymbiont *Rickettsia* also does not appear to be influencing population structure. In other insects, *Rickettsia* has been found to protect its host against heat shock (Brumin et al., 2011; Montllor et al., 2002) and pathogenic fungi (Łukasik et al., 2013), and could be adaptive in *A. permutata*. Further study is needed to make such a conclusion.

#### Insecticide resistance - adaptation to a highly modified landscape

Insecticide use is one of the most important stressors insects must overcome in human-altered landscapes (Douglas et al., 2020; Guedes et al., 2026; Hawkins et al., 2019; Serrão et al., 2022; Sowa et al., 2022; Wagner et al., 2021). Resistance signatures appear in all three clusters, but the vineyard-associated Clusters 1 and 3 carry additional ones—consistent with more intensive insecticide exposure in vineyards. Nonetheless, our findings show how sympatric populations exposed to the same insecticide regimes can evolve distinct adaptations for survival in human-altered environments, with implications for agricultural practice.

These inferences are exploratory. They rest on low-coverage data mapped to a congeneric reference (*A. alni*) and on the overlap of selection signatures with genes characterized in other insects, so they identify candidates for resistance rather than confirming it; a well-annotated *A. permutata* reference and higher-coverage resequencing could likely reveal further resistance-implicated genes missed here, and it is unlikely that every gene assigned to our five families of interest plays an adaptive role. We therefore treat genes that have been experimentally implicated in insecticide resistance elsewhere, and that fall within our genomic windows of interest, as quite plausibly contributing to resistance here, while genes assigned to these families but lacking such experimental support, we regard as primarily exploratory. Below, we discuss how insecticide regimes in our sampled localities may have driven selection in five gene families implicated in insecticide resistance, with functions—and potential management implications for vineyard *A. permutata*—that vary by gene family.

#### Carboxyl/cholinesterases

Both genetic Clusters 1 and 3 (primarily composed of vineyard individuals) show putative genetic sweeps in a chromosome 7 window containing a carboxylesterase gene (Supplemental Table S9), and all three populations show little structure in two windows containing carboxyl/cholinesterases, which hydrolyze organophosphates (OPs; Supplemental Table S10). Carboxylesterase amplification increases sequestration and hydrolysis of OPs (Bass et al., 2014; Field et al., 1988; Mouchès et al., 1986), which target acetylcholinesterase receptors (ace-1, Alout et al., 2007; Cheung et al., 2018). Given the history of OP usage in our study region (Table 4), we propose that this gene family could be broadly important to persistence alongside insecticide usage generally, while Cluster 1 and Cluster 3 harbor additional adaptations consistent with intensive insecticide usage in vineyards.

**Table 4.**
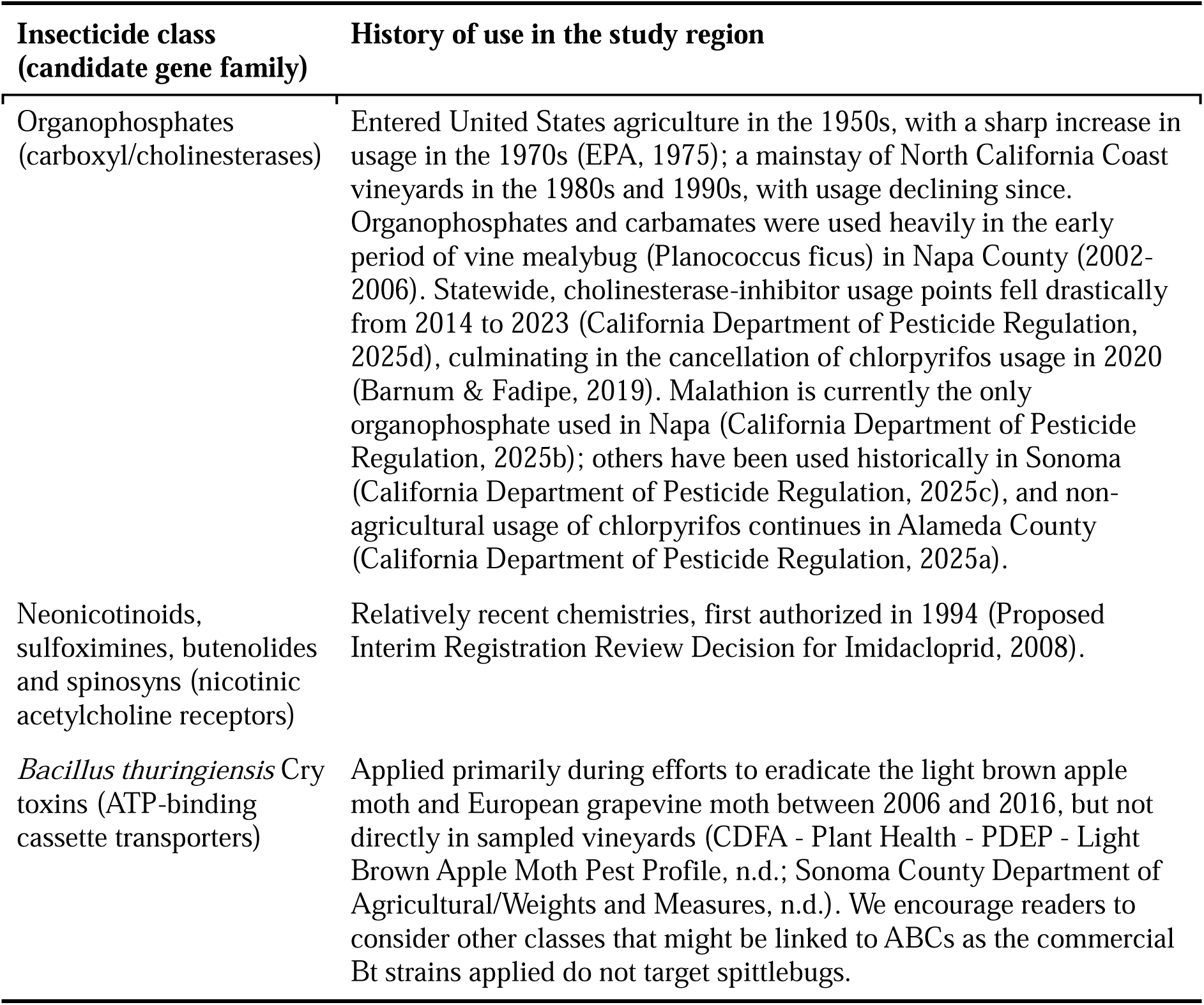
History of insecticide use in Napa, Sonoma and Alameda counties for the insecticide classes targeting the candidate resistance cal.

#### Cytochrome P450 Monooxygenases

Autosomal Cluster 3 was the only population with evidence of sweeps in the Cytochrome P450 monooxygenase gene family, for which overexpression, duplication and regulatory change drive resistance to DDT and neonicotinoids (Battlay et al., 2016; P. Daborn et al., 2001; Joußen et al., 2008; Pyke et al., 2004). We identify two specific genes in these swept regions which have been experimentally associated with insecticide resistance: CYP307a1 (Zhang et al., 2020; Etebari et al., 2018; Wu et al., 2018; Lv et al., 2026; Omoke et al., 2024; İnal et al., 2026) and CYP6a9 (Cucini et al., 2025; Fouda et al., 2025; Maitra et al., 1996). Given the varied history of insecticide applications in Central California (Table 4), and the wide variety of insecticides for which Cytochrome P450 monooxygenases might confer resistance, further research involving rearing *A. permutata* in the presence of insecticides and transcriptome sequencing is needed to determine specific insecticides for which Cluster 3 might have evolved increased resistance.

#### Nicotinic acetylcholine receptors

Nicotinic acetylcholine receptors (nAChR) are the direct target site for neonicotinoids, sulfoximines (sulfoxaflor), and butenolides (flupyradifurone). Mutations resulting in structural changes in nAChRs can change binding affinities, which are associated with insecticide resistance (Bass et al., 2011; Xu et al., 2022). Resistance to Spinosad (which has been used in California since 1994; Table 4) has been repeatedly mapped to a loss-of-function mutation in nAChRα6 (Perry et al., 2007, 2025; Zimmer et al., 2016), which has been swept in autosomal Cluster 2 (Supplemental Table S9). From the same family as nAChRα6, nAChRα5 was swept in autosomal Cluster 2, while nAChRα7, and nAChRβ1 (Supplemental Table S9), were swept in Cluster 1; and nAChRα8 showed little structure between Clusters 1 and 3 (Supplemental Table S10).

#### Additional Detoxification families

Two further gene families yielded resistance-associated signals. ATP-binding cassette (ABC) transporters act as midgut receptors for Crystal (Cry) toxins produced by Bacillus thuringiensis (Bt), and loss-of-function mutations confer robust resistance to Bt insecticides (Fabrick et al., 2022; Gahan et al., 2010; Park et al., 2014). We found low genetic structure in two ABC transporters between autosomal Clusters 1 and 3 (Supplemental Table S10) and sweeps of four ABC transporters across Clusters 2 and 3, which could reflect residual effects of recent Bt applications in the region (Table 4). Caution is merited in interpreting these results, as Bt toxins are generally not expected to be ingested by Hemiptera with piercing-sucking mouthparts and feeding behaviors (Chougule and Bonning, 2012), although low levels of uptake have been previously reported in insects with this feeding strategy (Burgio et al. 2007, 2011) or these genes could betide to natural Bt strains in the environment. UDP-glycosyltransferases (UGTs) glycosylate lipophilic toxins to raise their water solubility and speed excretion, and UGT overexpression is associated with resistance to a wide range of insecticides (Logan et al., 2024; Ma et al., 2021; Zhao et al., 2019). Here, one UGT-family gene showed low differentiation between Clusters 1 and 3 and another was swept in Cluster 3 (Supplemental Tables S9, S10); but because UGTs confer resistance so broadly, we cannot attribute these signals to insecticides specifically rather than some other selective pressure.

#### Future study and management implications

Testing the associations identified here will require rearing individuals in the presence of insecticides currently and historically applied in central California. Further work on how this cryptic structure and insecticide resistance interact with *X. fastidiosa* transmission could be valuable for stakeholders. Our demographic models (Table 3; Fig. 4) indicate ongoing gene flow between these populations, which may allow them to share resistance loci—another dynamic relevant to managing *X. fastidiosa*.

### Aphrophora taxonomy and species boundaries

#### The Vineyard Aphrophora is A. permutata

We approached this study with the expectation that vineyard *Aphrophora* would prove to be a cryptic species closely related to A. permutata, given the differing oviposition sites and adult migratory behaviors observed in vineyard and quasi-natural populations (Supplemental Note S1). Vineyard adults disappear during the prolonged, intense summer dry season; our limited observations at one locality suggest they may spend the summer on Douglas fir, though this needs broader confirmation. Wherever they go, they return to lay eggs in or on herbaceous vegetation—a complete departure from the quasi-natural life history, in which eggs are deposited in the needles of overstory pines and nymphs hatch the following spring and fall onto the understory. This vineyard life history requires changes in two complex behaviors: abandoning oviposition in conifer needles for the herbaceous understory, and replacing a short, one-way commute from understory to conifer canopy with a long-distance seasonal migration to summer hosts and back. Despite these differences, vineyard populations are conspecific with the quasi-natural Berkeley/Tilden population in which the detailed life history of *A. permutata* was studied (F_ST_ < 0.05).

We are unaware of comparably radical divergence in migration and oviposition behavior between geographic populations of the same species, made all the more remarkable here by their close proximity, all within a radius of about 40 km (Fig. 1C-D). In light of this extraordinary outcome, it would be informative to extend genome comparisons geographically and taxonomically to encompass widespread populations of *A. permutata* and closely related taxa.

#### Taxonomic implications across Aphrophora

The COI tree (Supplemental Fig. S6) is broadly consistent with *previous analyses of relationships in Aphrophora* (Doering 1941; Moore 1956; Hamilton 1982). All our samples are recovered in a polytomous clade along with *A. permutata*, *A. fulva* and *A. maculosa*. The seeming paraphyly we recover here is not surprising. Moore (1956) inferred these three species to be conspecific. While we make no formal taxonomic judgments on the validity of *A. fulva* and *A. maculosa* as separate species, we suspect that whole-genome analysis of putative individuals of these species may validate Moore’s (1956) judgement that they are synonymous.

The permutata group clade, described above, also includes several individuals identified as Aphrophora gelida (Supplemental Fig. S6), a species with very distinct male genitalia (Doering, 1941; Moore, 1956) and a transcontinental distribution, in contrast to the Rocky Mountains/Pacific-coastal range of the permutata group (Doering, 1941; Moore, 1956; Hamilton, 1982). Many of the *A. gelida* specimens included in the analysis were collected well outside the geographical range of the permutata group, making misidentification an unlikely explanation for their placement. These results suggest historical hybridization between *A. gelida* and members of the permutata group. Moore (1956) and Hamilton (1982) state that *A. gelida* is closely related to *A. permutata* and Hamilton places it in the same subgenus (*Plesiommata*). Combined with their overlapping distributions in the Canadian west and shared life histories living as adults on conifers (Hamilton 1982), this suggests that there may have been ample opportunity for introgression.

The present distinction in common names between *A. permutata* as the Douglas fir spittlebug and *A. fulva* as the Western pine spittlebug is clearly untenable, made all the more so by the evidence presented here that the species *permutata*, *fulva* and *maculosa* are indistinguishable by COI. Confirmation of this taxonomic finding will require examination of geographically diverse populations representing a much larger portion of the geographic range, stretching from Southern California north to Canada and east to the Rocky Mountains.

Perhaps the most interesting unresolved question, however, is how genetically similar, geographically adjacent populations maintain such different life histories, while facing similar anthropogenic pressures. *A. permutata* is widely distributed across North America from the Rocky Mountains west, raising opportunities to study its life cycle in geographically diverse settings. Is the life cycle in the Sonoma/Napa populations anomalous or widespread? Might the Tilden Park population, studied by Kelson (1964), be anomalous? Though Kelson’s (1964) study has not been replicated, in the earliest report on the pattern of occurrence of *A. permutata* nymphs, Ball (1901) reported that “Rocky Mountain region” nymphs have never been found “except where they were within a short distance of a pine tree,” but went on to say that, nonetheless, numerous individuals were found on plants in places apparently inaccessible to nymphs dropping from pines. He speculated that it appears “nearly certain that the adults must fly back to the [nymphal host] plants to deposit their eggs.” One hundred and twenty-five years later we are on the horns of the same dilemma. Do they or don’t they, and if so, where, in what circumstances, and why?

## Conclusion and Future Directions

We show that vineyard Aphrophora is one species with two cryptic, sympatric populations despite radically different life histories. The present study validates the original species identification of DeLong and Severin (1950) and demonstrates that *A. permutata* is untethered from the necessary presence of pines or other coniferous hosts in the immediate vicinity of vineyards. Unfortunately, this means that should *A. permutata* become a significant vector, it will not be amenable to one of the major approaches to control of *Homalodisca vitripennis*, the *X. fastidiosa* vector in Southern California vineyards. This sharpshooter leafhopper inhabits citrus as an alternative host and can be controlled in part by managing its presence in citrus orchards adjacent to grape plantings (Byrne & Redak, 2021). The alternative summer hosts of *A. permutata* appear to be far from vineyards and, so far, not identified. That said, if the summer hosts can be identified with certainty—Douglas fir being the leading candidate—managing *A. permutata* on those hosts could offer a control avenue analogous to the citrus-based suppression of *H. vitripennis*. Within the vineyards, Beal et al. (2021) have identified several preferred nymphal herbaceous hosts that might offer target opportunities for suppression. More broadly, this work provides evidence that sympatric populations can have distinct adaptations to anthropogenic pressures, such as insecticide use. This study further shows that endosymbiont infection dynamics warrant further investigation in adaptation to both urban and agricultural habitats. Priorities for future work include confirming the summer host(s), rearing experiments paired with transcriptome sequencing to test the candidate resistance loci identified here, and broader geographic and taxonomic sampling to resolve the status of *A. permutata*, *A. fulva*, and *A. maculosa*.

## Data availability

Raw reads have been uploaded to NCBI (in processing). The COI alignments, R markdown file and files used for statistical analyses have been uploaded to figshare (10.6084/m9.figshare.33812272).

## Acknowledgements

This work was funded by the California Department of Food and Agriculture Agreement Number 23-0382-000-SA to MKK and VT through the research support program of the California Pierce’s Disease/Glassy-Winged Sharpshooter Board. Sonoma vineyard specimens were supplied by Lucia Varela, Division of Agriculture and Natural Resources, University of California, Cooperative Extension, Santa Rosa. Napa vineyard specimens were supplied by Monica Cooper and Amielia Adams, Division of Agriculture and Natural Resources, University of California, Cooperative Extension, Napa. VT benefitted from discussions and vineyard field visits with these individuals and Dylan Beal and Cindy Kron. We also thank the growers who made their vineyards available for research, Jessica Ware for laboratory space at AMNH, and undergraduate students Priscilla Ramchand and Mark Mollo for help with DNA extractions. Computing was conducted through advanced research computing at Virginia Tech.

