## Supplementary Materials for "One species, two lives: cryptic population structure and life-history divergence in a vineyard spittlebug (*Aphrophora sp.*), a candidate vector for Pierce’s disease"

**Supplemental Figures**

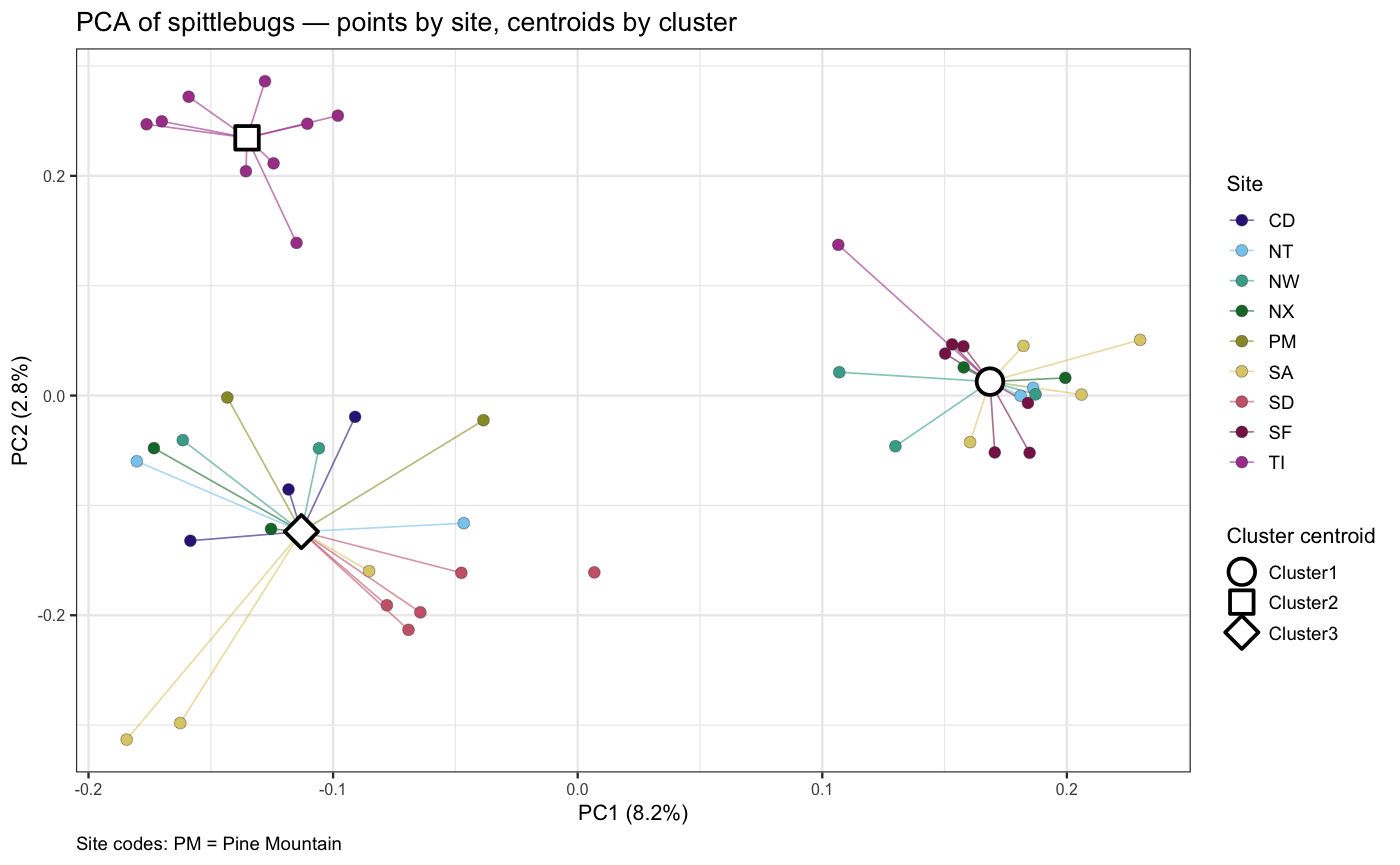

***Supplemental Fig. S1. Autosomal population structure by sampled site.***  *Principal Components Analysis (PCA) is estimated from nuclear SNPs. TI, CD and PM correspond to the three “quasi-natural” sampling sites*

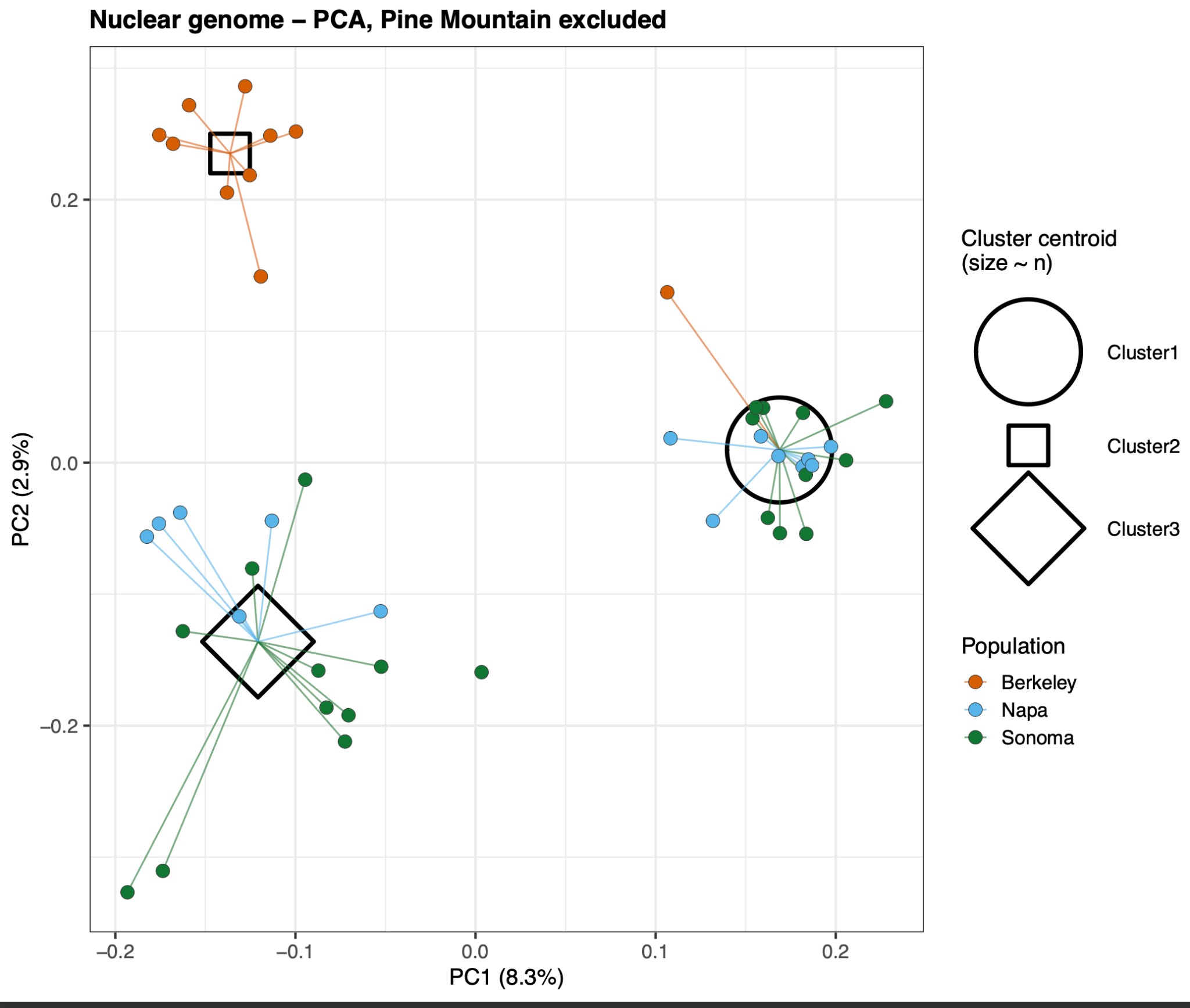

*Supplemental Fig. S2. Principal Components Analysis (PCA) from nuclear SNPs, excluding the two lowest coverage individuals from Pine Mountain. Individuals are coloured according to sampling locality (Berkeley, Napa, Sonoma and Pine Mountain), with the centroids for individuals assigned to Clusters 1, 2 and 3 in the autosomal PCA demarcated by a circle, square and diamond respectively.*

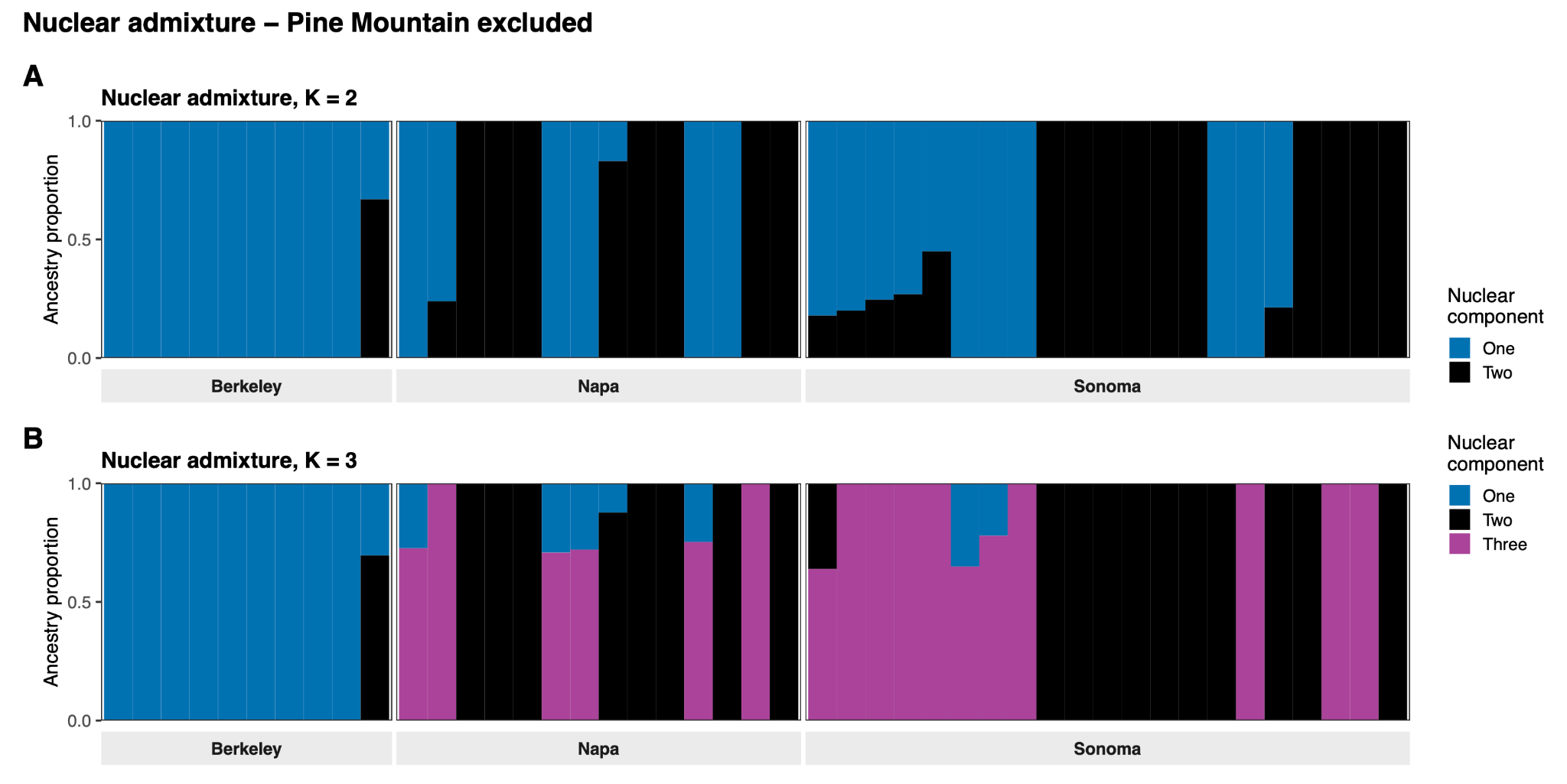

*Supplemental Fig. S3. Admixture components from nuclear SNPs at K=2 and K=3 model admixture, excluding the two lowest coverage individuals from Pine Mountain*

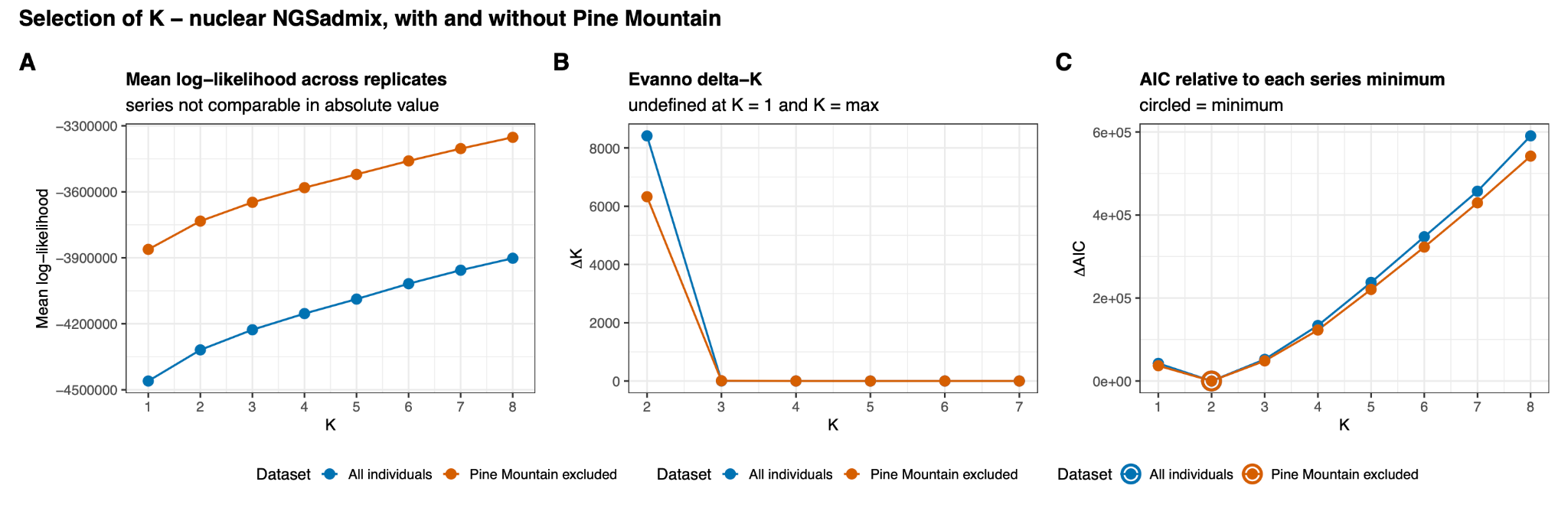

***Supplemental Fig. S4. Selection of K across nuclear admixture analyses.*** *A) Mean log-likelihood across the ten replicates of K. The proportion of replicates within 2 log-likelihood of the best run are shown. B) Evanno ΔK across K=2 to K=7. C) AIC across replicates of K. The lowest value is encircled. Analyses are shown with (blue) and without (orange) the two lowest coverage individuals from Pine Mountain.*

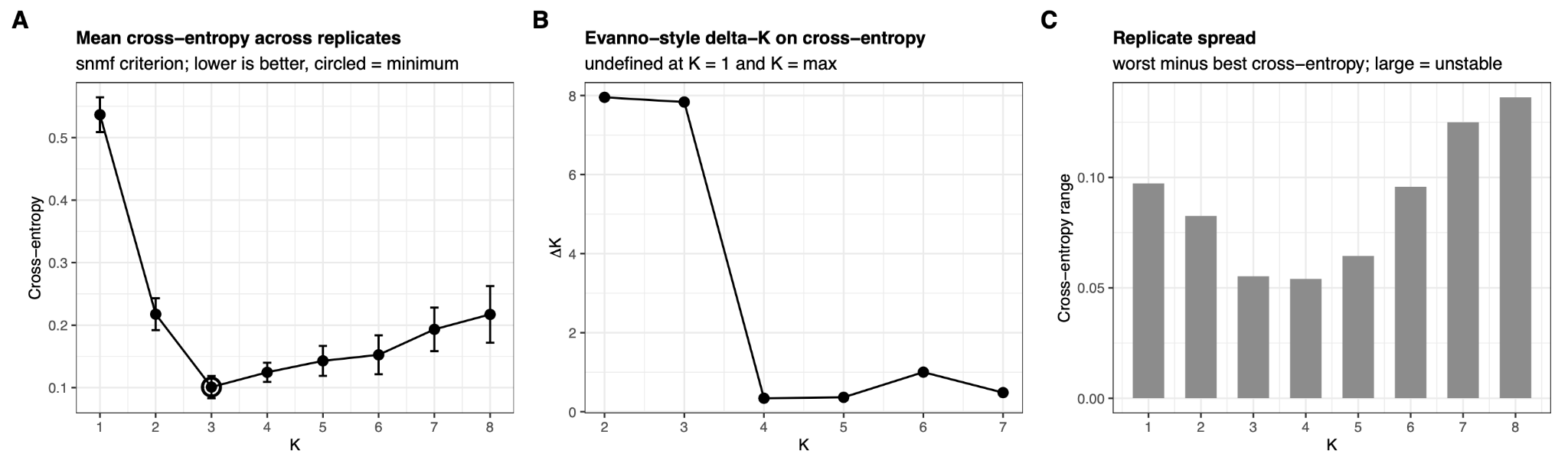

***Supplemental Fig. S5. Sparse non-negative matrix factorization fit at different levels of K.*** *A) Mean cross-entropy across replicates with the smallest value circled. B) Evanno-style ΔK on cross-entropy. Note this value is undefined at K=1 and K=max. C) Spread of cross-entropy across replicates. Larger values indicate greater model instability. Ten replicates were conducted for each level of K.*

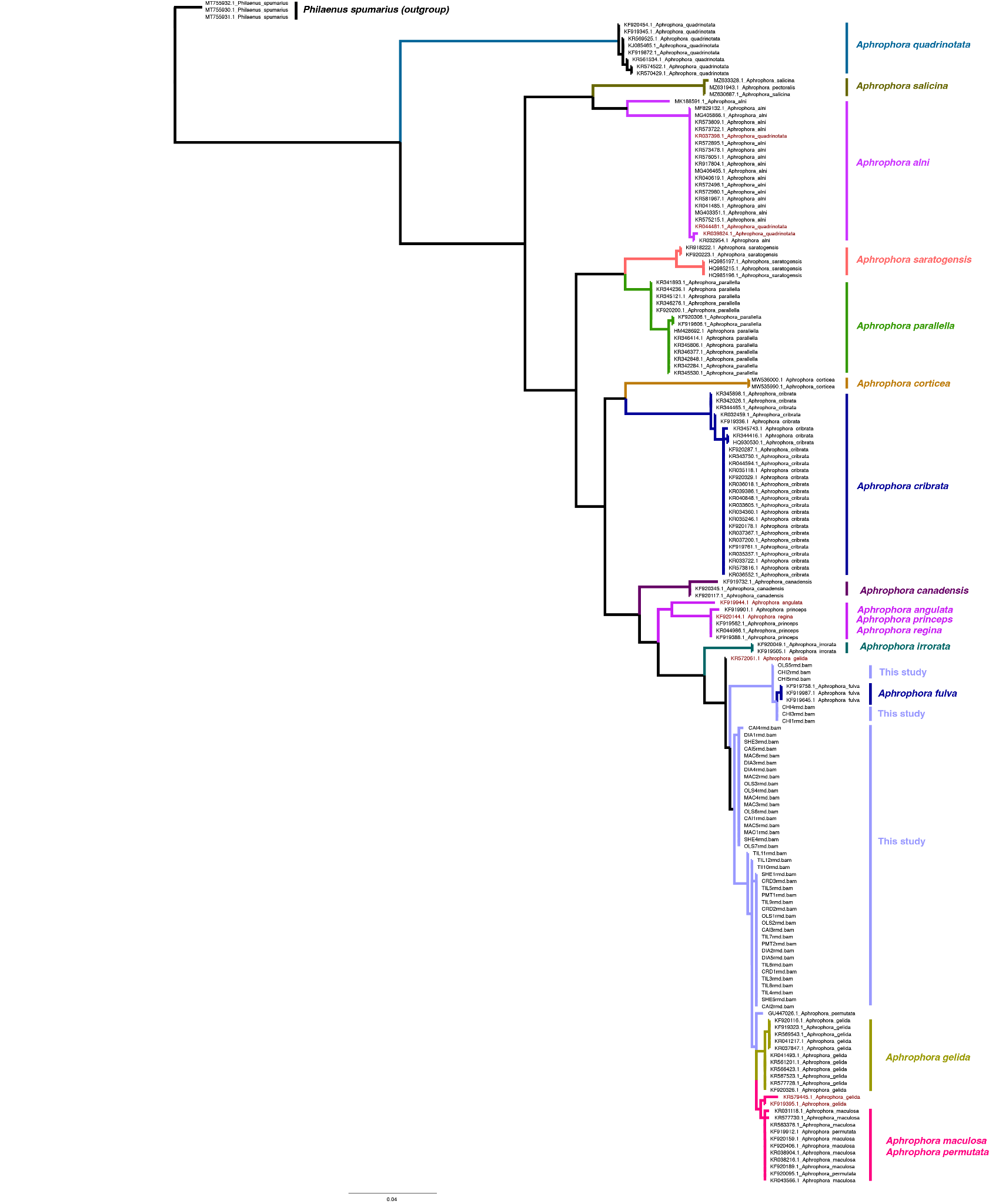

***Supplemental Fig. S6. COI tree for Aphrophora permutata and closely related species.*** All the specimens analyzed for this study fall in a clade of closely related individuals that include specimens previously identified as A. permutata, *A. fulva*, *A. maculosa*, and *A. gelida*. For the first three species (the *permutata* group) and our specimens, this close relationship is consistent with the hypothesis that they are representative of a single species. The embedded *A. gelida* specimens appear to represent a case of mitochondrial capture between closely related species (see text). The *A. quadrinotata* specimens in the *A. alni* clade are probably misidentifications (these species are superficially close in appearance). The *A. regina* specimen in the *A. princeps* clade is consistent with the close relationship between these species (Hamilton 1982).

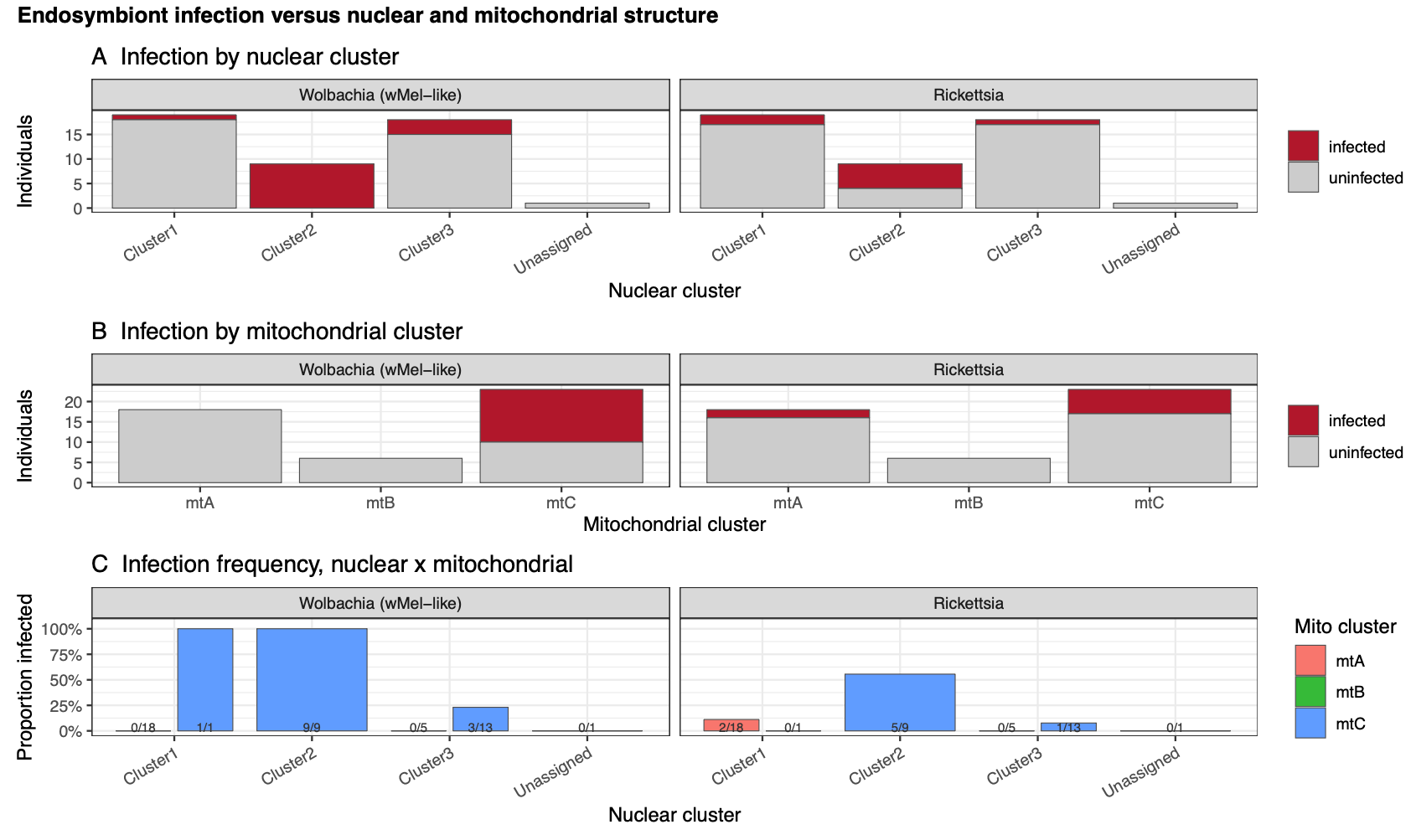

***Supplemental Fig. S7. Endosymbiont infection by nuclear and mitochondrial lineage.*** *A) Proportion of individuals infected by Wolbachia and Rickettsia across our three identified autosomal clusters. B) Proportion of individuals infected by Wolbachia and Rickettsia across our three identified mitochondrial lineages. C) Wolbachia and Rickettsia infection by both mitochondrial lineage, and autosomal cluster.*

**Supplemental Tables**

**Table S1. Specimens of Aphrophora permutata included in this study, with collection details and mean sequencing coverage. Site codes follow the anonymised scheme used throughout the manuscript.**

| *Site* | *Region* | *Collection Date* | *Preservation method* | *Life stage* | *Specimen* | *Host plant*  *If known* | *Mean coverage (×)* |
| --- | --- | --- | --- | --- | --- | --- | --- |
| SF | Sonoma | May 9th 2019 | Ethanol | 5th instar nymph | One |  | 1.62 |
| SF | Sonoma | May 9th 2019 | Ethanol | 5th instar nymph | Two |  | 1.15 |
| SF | Sonoma | May 9th 2019 | Ethanol | 5th instar nymph | Three |  | 1.52 |
| SF | Sonoma | May 9th 2019 | Ethanol | 5th instar nymph | Four |  | 1.54 |
| SF | Sonoma | May 9th 2019 | Ethanol | 5th instar nymph | Five |  | 1.21 |
| SF | Sonoma | May 9th 2019 | Ethanol | 5th instar nymph | Six |  | 1.47 |
| SD | Sonoma | May 10th 2019 | Ethanol | 5th instar nymph | One |  | 1.39 |
| SD | Sonoma | May 10th 2019 | Ethanol | 5th instar nymph | Two |  | 1.60 |
| SD | Sonoma | May 10th 2019 | Ethanol | 5th instar nymph | Three |  | 1.22 |
| SD | Sonoma | May 10th 2019 | Ethanol | 5th instar nymph | Four |  | 1.38 |
| SD | Sonoma | May 10th 2019 | Ethanol | 5th instar nymph | Five |  | 1.38 |
| CD | Sonoma | June 3 2024 | Ethanol | Adult | One | *Pseudotsuga menziesii* | 1.59 |
| CD | Sonoma | June 3 2024 | Ethanol | Adult | Two | *Pseudotsuga menziesii* | 1.71 |
| CD | Sonoma | June 3 2024 | Ethanol | Adult | Three | *Pseudotsuga menziesii* | 1.63 |
| SA | Sonoma | May 2019 | Ethanol | Adult | One |  | 1.45 |
| SA | Sonoma | May 15 2019 | Ethanol | Adult | Two |  | 1.60 |
| SA | Sonoma | May 13 2019 | Ethanol | Adult | Three |  | 1.43 |
| SA | Sonoma | May 15 2019 | Ethanol | Adult | Four |  | 1.40 |
| SA | Sonoma | May 19 2019 | Ethanol | Adult | Five |  | 1.80 |
| SA | Sonoma | May 19 2019 | Ethanol | Adult | Six |  | 1.81 |
| SA | Sonoma | May 19 2019 | Ethanol | Adult | Seven |  | 1.45 |
| TI | Berkeley, California | June 14th 2024 | Ethanol | Adult | Three | *Pseudotsuga menziesii* | 1.64 |
| TI | Berkeley, California | June 14th 2024 | Ethanol | Adult | Four | *Pseudotsuga menziesii* | 1.54 |
| TI | Berkeley, California | June 11th 2024 | Ethanol | Adult | Five | *Pinus radiata* | 1.58 |
| TI | Berkeley, California | June 11th 2024 | Ethanol | Adult | Six | *Pinus radiata* | 1.64 |
| TI | Berkeley, California | June 11th 2024 | Ethanol | Adult | Seven |  | 1.28 |
| TI | Berkeley, California | June 11th 2024 | Ethanol | Adult | Eight |  | 1.44 |
| TI | Berkeley, California | June 11th 2024 | Ethanol | Adult | Nine |  | 1.89 |
| TI | Berkeley, California | June 11th 2024 | Ethanol | Adult | Ten |  | 1.41 |
| TI | Berkeley, California | June 11th 2024 | Ethanol | Adult | Eleven |  | 1.44 |
| TI | Berkeley, California | June 11th 2024 | Ethanol | Adult | Twelve |  | 1.73 |
| PM | Marin County | May 26th 2022 | Ethanol | Adult | One | *Pseudotsuga menziesii* | 0.49 |
| PM | Marin County | May 26th 2022 | Dried from ethanol | Adult | Two | *Pinus radiata* | 0.69 |
| NT | Napa | April 5th 2019 | Ethanol | Early instar nymph | One | *Chlorogalum pomeridanum* | 1.62 |
| NT | Napa | April 5th 2019 | Ethanol | Early instar nymph | Two | *Chlorogalum pomeridanum* | 1.67 |
| NT | Napa | April 5th 2019 | Ethanol | Early instar nymph | Three | *Chlorogalum pomeridanum* | 1.51 |
| NT | Napa | April 5th 2019 | Ethanol | Early instar nymph | Four | *Chlorogalum pomeridanum* | 1.57 |
| NT | Napa | April 5th 2019 | Ethanol | Early instar nymph | Five | *Chlorogalum pomeridanum* | 1.48 |
| NX | Napa | May 2019 | Ethanol | Adult | One |  | 1.00 |
| NX | Napa | May 2019 | Ethanol | Adult | Three |  | 0.63 |
| NX | Napa | May 2019 | Dried from ethanol | Adult | Four |  | 1.53 |
| NX | Napa | May 2019 | Dried from ethanol | Adult | Five |  | 1.25 |
| NW | Napa | May 2019 | Ethanol | Adult | One |  | 1.55 |
| NW | Napa | April 11th 2019 | Ethanol | 5th instar | Two |  | 1.39 |
| NW | Napa | April 11th 2019 | Ethanol | 5th instar | Three |  | 1.79 |
| NW | Napa | April 11th 2019 | Ethanol | 5th instar | Four |  | 2.32 |
| NW | Napa | April 11th 2019 | Pinned | 5th instar | Five |  | 1.41 |
| For cross reference, the Sonoma County sites correspond to the sites coded as SA, SD, and SF in Beal et al. 2021 (Table S1), site codes we replicate here. One Napa County site corresponds to their site coded as NT. The other two Napa County sites we have arbitrarily assigned codes NW and NX, which do not overlap those of Beal et al. | | | | | | |  |

| ***Supplemental Table S2. Flags and options used in analyses.*** | |
| --- | --- |
| Analysis | Added flags and options |
| Genotype likelihood calculation (ANGSD) | -remove_bads 1 -uniqueOnly 1 -only_proper_pairs 1 -C 50 -baq 1 -minMapQ 30 -minQ 30 -minInd 30 -doSnpStat 1 -doHWE 1 -sb_pval 1e-6 -hwe_pval 1e-6 -hetbias_pval 1e-6 -doMajorMinor 1 -skipTriallelic 1 -doMaf 2 -doPost 1 -minMaf 0.05 -snp_pval 1e-6 -doGeno 8 -doGlf 2 |
| Linkage identification and pruning (NGSLD) | --field_dist 3 --max_kb_dist 50000 --field_weight7 --min_weight 0.1 |
| Mitochondrial sequence alignment (MAFT) | --auto |

Table S3. Publicly available sequences used in COI analyses.

| **Accession number** | **Species** |
| --- | --- |
| GU447026.1 | *Aphrophora permutata* |
| HM428692.1 | *Aphrophora parallella* |
| HQ930530.1 | *Aphrophora cribrata* |
| HQ985196.1 | *Aphrophora saratogensis* |
| HQ985197.1 | *Aphrophora saratogensis* |
| HQ985215.1 | Aphrophora saratogensis |
| KF919323.1 | Aphrophora gelida |
| KF919336.1 | Aphrophora cribrata |
| KF919345.1 | Aphrophora quadrinotata |
| KF919388.1 | Aphrophora princeps |
| KF919395.1 | Aphrophora gelida |
| KF919505.1 | Aphrophora irrorata |
| KF919562.1 | Aphrophora princeps |
| KF919606.1 | Aphrophora parallella |
| KF919645.1 | Aphrophora fulva |
| KF919732.1 | Aphrophora canadensis |
| KF919758.1 | Aphrophora fulva |
| KF919761.1 | Aphrophora cribrata |
| KF919872.1 | Aphrophora quadrinotata |
| KF919901.1 | Aphrophora princeps |
| KF919912.1 | Aphrophora permutata |
| KF919944.1 | Aphrophora angulata |
| KF919987.1 | Aphrophora fulva |
| KF920049.1 | Aphrophora irrorata |
| KF920095.1 | Aphrophora permutata |
| KF920116.1 | Aphrophora gelida |
| KF920117.1 | Aphrophora canadensis |
| KF920144.1 | Aphrophora regina |
| KF920159.1 | Aphrophora maculosa |
| KF920178.1 | Aphrophora cribrata |
| KF920189.1 | Aphrophora maculosa |
| KF920200.1 | Aphrophora parallella |
| KF920223.1 | Aphrophora saratogensis |
| KF920287.1 | Aphrophora cribrata |
| KF920306.1 | Aphrophora parallella |
| KF920326.1 | Aphrophora gelida |
| KF920329.1 | Aphrophora cribrata |
| KF920345.1 | Aphrophora canadensis |
| KF920406.1 | Aphrophora maculosa |
| KF920454.1 | Aphrophora quadrinotata |
| KJ085465.1 | Aphrophora quadrinotata |
| KR031118.1 | Aphrophora maculosa |
| KR032459.1 | Aphrophora cribrata |
| KR032954.1 | Aphrophora alni |
| KR033605.1 | Aphrophora cribrata |
| KR033722.1 | Aphrophora cribrata |
| KR034360.1 | Aphrophora cribrata |
| KR035118.1 | Aphrophora cribrata |
| KR035246.1 | Aphrophora cribrata |
| KR035357.1 | Aphrophora cribrata |
| KR036018.1 | Aphrophora cribrata |
| KR036552.1 | Aphrophora cribrata |
| KR037200.1 | Aphrophora cribrata |
| KR037367.1 | Aphrophora cribrata |
| KR037398.1 | Aphrophora quadrinotata |
| KR037847.1 | Aphrophora gelida |
| KR038216.1 | Aphrophora maculosa |
| KR038904.1 | Aphrophora maculosa |
| KR039386.1 | Aphrophora cribrata |
| KR039824.1 | Aphrophora quadrinotata |
| KR040619.1 | Aphrophora alni |
| KR040848.1 | Aphrophora cribrata |
| KR041217.1 | Aphrophora gelida |
| KR041485.1 | Aphrophora alni |
| KR041493.1 | Aphrophora gelida |
| KR043566.1 | Aphrophora maculosa |
| KR044481.1 | Aphrophora quadrinotata |
| KR044594.1 | Aphrophora cribrata |
| KR044986.1 | Aphrophora princeps |
| KR341893.1 | Aphrophora parallella |
| KR342026.1 | Aphrophora cribrata |
| KR342284.1 | Aphrophora parallella |
| KR342848.1 | Aphrophora parallella |
| KR343750.1 | Aphrophora cribrata |
| KR344236.1 | Aphrophora parallella |
| KR344416.1 | Aphrophora cribrata |
| KR344465.1 | Aphrophora cribrata |
| KR345121.1 | Aphrophora parallella |
| KR345530.1 | Aphrophora parallella |
| KR345743.1 | Aphrophora cribrata |
| KR345806.1 | Aphrophora parallella |
| KR345898.1 | Aphrophora cribrata |
| KR346276.1 | Aphrophora parallella |
| KR346377.1 | Aphrophora parallella |
| KR346414.1 | Aphrophora parallella |
| KR561201.1 | Aphrophora gelida |
| KR561534.1 | Aphrophora quadrinotata |
| KR563376.1 | Aphrophora maculosa |
| KR566423.1 | Aphrophora gelida |
| KR567523.1 | Aphrophora gelida |
| KR569525.1 | Aphrophora quadrinotata |
| KR569543.1 | Aphrophora gelida |
| KR570429.1 | Aphrophora quadrinotata |
| KR572061.1 | Aphrophora gelida |
| KR572496.1 | Aphrophora alni |
| KR572895.1 | Aphrophora alni |
| KR572980.1 | Aphrophora alni |
| KR573478.1 | Aphrophora alni |
| KR573722.1 | Aphrophora alni |
| KR573809.1 | Aphrophora alni |
| KR573816.1 | Aphrophora cribrata |
| KR574522.1 | Aphrophora quadrinotata |
| KR575215.1 | Aphrophora alni |
| KR576051.1 | Aphrophora alni |
| KR577728.1 | Aphrophora gelida |
| KR577730.1 | Aphrophora maculosa |
| KR579445.1 | Aphrophora gelida |
| KR581967.1 | Aphrophora alni |
| KR917804.1 | Aphrophora alni |
| KR918222.1 | Aphrophora saratogensis |
| MF829132.1 | Aphrophora alni |
| MG403351.1 | Aphrophora alni |
| MG405866.1 | Aphrophora alni |
| MG406465.1 | Aphrophora alni |
| MK188591.1 | Aphrophora alni |
| MT755930.1 | Philaenus spumarius |
| MT755931.1 | Philaenus spumarius |
| MT755932.1 | Philaenus spumarius |
| MW535990.1 | Aphrophora corticea |
| MW536000.1 | Aphrophora corticea |
| MZ630687.1 | Aphrophora salicina |
| MZ631943.1 | Aphrophora pectoralis |
| MZ633328.1 | Aphrophora salicina |

**Table S4.** Gene families screened for within outlier and conserved genomic windows of *Aphrophora permutata*, with the references used to define each family.

| Gene family | Abbreviation | Reference(s) |
| --- | --- | --- |
| ***Chemosensory: ligand capture*** | | |
| Odorant-binding proteins | OBP | Vogt & Riddiford (1981); Pelosi et al. (2018) |
| Chemosensory proteins | CSP | Pelosi et al. (2018) |
| ***Chemosensory: membrane transfer*** | | |
| Sensory neuron membrane proteins | SNMP | Benton et al. (2007) |
| ***Chemosensory: signal transduction*** | | |
| Odorant receptors | OR | Clyne et al. (1999); Vosshall et al. (1999) |
| Gustatory receptors | GR | Clyne et al. (2000); Scott et al. (2001) |
| Ionotropic receptors | IR | Benton et al. (2009); Croset et al. (2010) |
| ***Insecticide resistance: metabolic detoxification*** | | |
| Cytochrome P450 monooxygenases | P450 | Nauen et al. (2022) |
| Carboxyl/cholinesterases | — | Oakeshott et al. (2005) |
| UDP-glycosyltransferases | UGT | Zhou et al. (2019) |
| ATP-binding cassette transporters | ABC | Dermauw & Van Leeuwen (2014) |
| Glutathione S-transferases | — | Enayati et al. (2005) |
| ***Insecticide resistance: target site*** | | |
| Nicotinic acetylcholine receptors | nAChR | Millar & Denholm (2007) |
| Voltage-gated sodium channels | — | Dong et al. (2014) |
| Acetylcholinesterases | — | Weill et al. (2003) |
| γ-Aminobutyric acid (GABA) receptors | — | Ffrench-Constant et al. (2000) |
| Ryanodine receptors | — | Sattelle et al. (2008) |
| Glutamate-gated chloride channels | — | Wolstenholme (2012) |
| Chitin synthases | — | Douris et al. (2016) |

Families were identified among genes overlapping outlier and conserved windows using the eggNOG 5.0 functional annotation of the *Aphrophora alni* gene set (Huerta-Cepas et al., 2019; Cantalapiedra et al., 2021), with orthology restricted to Insecta.

— indicates that no abbreviation for the family is used in the main text.

**Table S5.** Reference genomes used to screen *Aphrophora permutata* resequencing reads for bacterial endosymbionts.

| Taxon | Supergroup | Strain | Reference |
| --- | --- | --- | --- |
| ***Endosymbiont reference panel*** | | | |
| *Wolbachia* sp. | A | wMel | Wu et al. (2004) |
| *Wolbachia* sp. | A | wRi | Klasson et al. (2009) |
| *Wolbachia* sp. | B | wAlbB | Klasson et al. (2008) |
| *Wolbachia* sp. | F | wCle | Sinha et al. (2019) |
| Cardinium hertigii | — | — | Penz et al. (2012) |
| *Rickettsia bellii* | — | — | Ogata et al. (2006) |
| *Spiroplasma poulsonii* | — | — | Masson et al. (2018) |
| *Arsenophonus nasoniae* | — | — | Wilkes et al. (2010) |
| ***Host reference*** | | | |
| *Aphrophora alni*^a^ | — | — | Griffiths et al. (2024) |

^a^ Included in the panel to distinguish reads deriving from *Wolbachia* sequence inserted in the host nuclear genome from reads deriving from a live infection (Dunning Hotopp et al., 2007).

Reads were mapped to the panel with BWA-MEM (Li & Durbin, 2009), retaining paired alignments with a mapping quality of at least 30. An individual was scored as infected at a coverage breadth greater than 0.30 and a Gini coefficient below 0.60.

**Table S6.** Thermal indices calculated for each of the nine sampling localities from Open-Meteo daily reanalysis of 2 m air temperature, 1995–2024. The activity season is the May–October adult flight period of *Aphrophora permutata*. These indices were tested against *Wolbachia* and *Rickettsia* infection status as described in the Materials and Methods.

| Index | Definition | Averaging window |
| --- | --- | --- |
| ***Mean temperatures*** | | |
| Annual average temperature | Mean of daily mean temperature | Calendar year, averaged across 1995–2024 |
| Activity-season average temperature | Mean of daily mean temperature | May–October, averaged across 1995–2024 |
| Average low temperature | Mean of daily minimum temperature | Activity season, averaged across 1995–2024 |
| ***Peak temperatures*** | | |
| Average peak temperature | Mean of daily maximum temperature | Activity season, averaged across 1995–2024 |
| Absolute peak temperature | Highest daily maximum temperature recorded | 1995–2024 |
| Yearly maximum average | Mean across years of each year's highest daily maximum temperature | Calendar year, averaged across 1995–2024 |
| ***Heat-day frequencies*** | | |
| Days exceeding 30 °C | Number of days with a daily maximum above 30 °C | Activity season, mean days per year |
| Days exceeding 32 °C | Number of days with a daily maximum above 32 °C | Activity season, mean days per year |
| Days exceeding 35 °C | Number of days with a daily maximum above 35 °C | Activity season, mean days per year |
| ***Variability*** | | |
| Mean daily temperature fluctuation | Mean of the daily maximum minus the daily minimum temperature | Activity season, averaged across 1995–2024 |

**Table S7.** Tests of association between per-individual sequencing depth and nuclear population structure in *Aphrophora* (n = 47).

**(a) Sequencing depth by genetic cluster**

| **Genetic cluster** | **n** | **Mean depth (×)** | **SD** | **Range (×)** |
| --- | --- | --- | --- | --- |
| Cluster 1 | 19 | 1.501 | 0.324 | 0.626 – 2.321 |
| Cluster 2 | 9 | 1.571 | 0.181 | 1.285 – 1.887 |
| Cluster 3 | 18 | 1.375 | 0.348 | 0.490 – 1.805 |
| Unassigned | 1 | 1.391 | — | — |

Kruskal–Wallis H = 2.327, df = 2, P = 0.312, ε² = 0.008. One-way ANOVA F = 1.386, P = 0.261. No pairwise contrast was significant (Mann–Whitney, BH-corrected: all q = 0.403). Restricting to the main sequencing batch gave the same result (H = 2.280, P = 0.320).

**(b) Rank correlations between depth and ordination and admixture scores**

| **Association tested** | **K** | **ρ** | **P** | **q (BH)** |
| --- | --- | --- | --- | --- |
| Depth ~ PC2 | — | 0.185 | 0.213 | 0.958 |
| Depth ~ max(Q) — assignment strength | 3 | 0.169 | 0.256 | 0.958 |
| Depth ~ entropy(Q) — apparent admixture | 3 | −0.169 | 0.256 | 0.958 |
| Depth ~ PC3 | — | 0.115 | 0.440 | 0.958 |
| Depth ~ ancestry component 3 | 3 | −0.102 | 0.496 | 0.958 |
| Depth ~ ancestry component 1 | 3 | 0.063 | 0.673 | 0.958 |
| Depth ~ max(Q) — assignment strength | 2 | 0.063 | 0.675 | 0.958 |
| Depth ~ entropy(Q) — apparent admixture | 2 | −0.063 | 0.675 | 0.958 |
| Depth ~ ancestry component 2 | 3 | −0.046 | 0.758 | 0.958 |
| Depth ~ PC1 | — | 0.012 | 0.935 | 0.958 |
| Depth ~ ancestry component 1 | 2 | −0.008 | 0.958 | 0.958 |
| Depth ~ ancestry component 2 | 2 | 0.008 | 0.958 | 0.958 |

Spearman rank correlations ρ(), ordered by P. The q column gives Benjamini–Hochberg values across all twelve tests. No test was significant at q < 0.05. Correlations with principal components are given as absolute values, the sign of an eigenvector being arbitrary.

Table S8. Demographic model comparison for the Cluster3–Cluster1 joint site frequency spectrum, projected to 16 × 18 haploid genomes. Models were fitted in GADMA2 with the moments engine (50 genetic-algorithm repeats each). AIC and CLAIC are reported for completeness; neither was used for selection (see notes).

| Model | Description | *k* | ln *L* | AIC | ΔAIC | CLAIC | ΔCLAIC | Penalty |
| --- | --- | --- | --- | --- | --- | --- | --- | --- |
| M3^a^ | Split, symmetric gene flow | 7 | −1,257.45 | 2,528.91 | 0.00 | 2,515.01 | 0.02 | 0.11 |
| M2 | Split, asymmetric gene flow | 8 | −1,257.40 | 2,530.79 | 1.89 | 2,514.99 | 0.00 | 0.19 |
| M1 | Split, no gene flow (null) | 4 | −3,279.55 | 6,567.09 | 4,038.19 | — | — | — |

*k* = number of free parameters. Penalty = CLAIC − (−2 ln *L*), the estimated 2 tr(*JH*⁻¹) term.

^a^ Selected model. M2 attained the marginally lower CLAIC, but the 0.02-unit difference is far below the ~2 units conventionally taken as a separation, and the two fits differ by 0.05 ln *L* units. M3 was selected as the more parsimonious of two indistinguishable models, consistent with its lower AIC.

**Table S9.** Insecticide-resistance candidate genes falling within a population branch excess (PBE) outlier window in at least one genetic cluster of *Aphrophora* *permutata.*

Genes are grouped by gene family. A gene is listed when it overlaps a 50 kb window in the upper 1% of the PBE distribution for that cluster Seventeen of 330 screened insecticide resistance gene candidates qualify. Annotations are from eggNOG-mapper; “—” denotes no assigned gene symbol.

| **Gene** | **Gene ID** | **Annotation** | **Chr^a^** | **Cluster(s)^b^** |
| --- | --- | --- | --- | --- |
| **Nicotinic acetylcholine receptors** | | | | |
| *nAChRa5* | ENSVPRG00000009328 | Neurotransmitter-gated ion channel | 3 | 2 |
| *nAChRa6* | ENSVPRG00000010379 | Acetylcholine-gated cation channel | 5 | 2 |
| *nAChRa7* | ENSVPRG00000009810 | Neurotransmitter-gated ion channel | 5 | 1 |
| *nAChRb1* | ENSVPRG00000012368 | Neurotransmitter-gated ion channel | 5 | 1 |
| — | ENSVPRG00000014997 | Ligand-gated ion channel (Neur_chan_LBD, Neur_chan_memb) | 4 | 2 |
| —^a^ | ENSVPRG00000015879 | Farnesoic acid O-methyltransferase / Neur_chan_LBD | 4 | 2 |
| **Cytochrome P450 monooxygenases** | | | | |
| *CYP307A1* | ENSVPRG00000008297 | Cytochrome P450 | 3 | 3 |
| *Cyp6a9* | ENSVPRG00000004142 | Cytochrome P450 | 2 | 3 |
| — | ENSVPRG00000011725 | Cytochrome P450 | 3 | 3 |
| — | ENSVPRG00000008433 | Cytochrome P450 | 6 | 3 |
| **Carboxyl/cholinesterases** | | | | |
| — | ENSVPRG00000006002 | Carboxylesterase | 7 | 1, 3 |
| **ATP-binding cassette transporters**  —  Phase III efflux of xenobiotics and their conjugates; also implicated in Bt toxin receptor function. | | | | |
| — | ENSVPRG00000004358 | Transmembrane transporter (ABC_membrane, ABC_tran) | 13 | 2 |
| — | ENSVPRG00000014140 | ABC-2 family transporter | X | 3 |
| — | ENSVPRG00000013950 | Multidrug resistance-associated protein (MRP-like) | 8 | 3 |
| — | ENSVPRG00000015553 | ABC transporter, transmembrane region | 8 | 3 |
| **UDP-glycosyltransferases** | | | | |
| *Ugt*^c^ | ENSVPRG00000013626 | UDP-glucose:glycoprotein glucosyltransferase | 8 | 2 |
| — | ENSVPRG00000010950 | Glycosyltransferase family 28 | 3 | 3 |

^a^ chromosome

^b^ Chimeric annotation: carries both Neur_chan_LBD and Methyltransf_FA domains and is described as a farnesoic acid O-methyltransferase. Family assignment is unreliable.

^c^ Carries the preferred name Ugt, but its PFAM assignment is UDP-g_GGTase (UDP-glucose:glycoprotein glucosyltransferase, an endoplasmic-reticulum folding quality-control enzyme), not a detoxification UGT. Listed for transparency and excluded from family-level inference.

Chromosome X is the sex chromosome; autosomes are numbered 1–14. Unplaced scaffolds and the mitochondrial genome were excluded from all analyses.

**Table S10.** Insecticide-resistance candidate genes in the most conserved windows of *Aphrophora permutata*

A window qualifies for a given comparison when its FST falls in the lowest 1% of that comparison's distribution (cutoffs 0.0338, 0.0269 and 0.0241 for Cluster 1–2, 1–3 and 2–3 respectively, of 27,468 windows). Panel (a) requires all three comparisons; panel (b) requires Cluster 1–3 and excludes windows already in panel (a).

(a) Undifferentiated across all three clusters — 51 windows, 40 genes, 3 pesticide resistance candidates.

| **Gene** | **Gene ID** | | **Annotation** | **Chr.** | **Window (bp)** |  | **FST range** |
| --- | --- | --- | --- | --- | --- | --- | --- |
|  | | **Carboxyl/cholinesterases** | | | | | |
| *Gli* | ENSVPRG00000006890 | | Carboxylesterase | 2 | 18,070,000–18,120,000 |  | 0.0200–0.0304 |
| — | ENSVPRG00000002730^*^ | | Alpha/beta hydrolase fold | 9 | 51,930,000–51,980,000 |  | 0.0101–0.0141 |
| — | ENSVPRG00000002767^(^ | | Alpha/beta hydrolase fold | 9 | 51,930,000–51,980,000 |  | 0.0101–0.0141 |

(b) Undifferentiated between Clusters 1 and 3 only — 224 windows, 155 genes, 5 pesticide resistance candidates

| **Gene** | **Gene ID** | **Annotation** | **Chr.** | **Window (bp)** | **FST range** |
| --- | --- | --- | --- | --- | --- |
| **Nicotinic acetylcholine receptors** | | | | | |
| *nAChRa8* | ENSVPRG00000010897 | Neurotransmitter-gated ion channel | X | 171,130,000–171,180,000 | 0.0240–0.0488 |
| — | ENSVPRG00000007120 | Neurotransmitter-gated ion channel | 12 | 37,520,000–37,570,000 | 0.0265–0.0575 |
| **ATP-binding cassette transporters** | | | | | |
| — | ENSVPRG00000003223 | Multidrug resistance protein | 11 | 48,550,000–48,600,000 | 0.0240–0.0444 |
| — | ENSVPRG00000006913 | ATP-binding cassette domain | 13 | 31,720,000–31,770,000 | 0.0240–0.0388 |
| **UDP-glycosyltransferases** | | | | | |
| — | ENSVPRG00000003656 | transferase activity, transferring hexosyl group | 10 | 44,280,000–44,330,000 | 0.0233–0.0934 |

^*^ ENSVPRG00000002730 and ENSVPRG00000002767 are adjacent paralogues in a single 50 kb window and are not independent observations.

| **Table S11.** Breadth, mean depth, and the standard Gini coefficient of sequencing coverage in individuals flagged as carriers of endosymbionts. | | | | |
| --- | --- | --- | --- | --- |
|  | **n infected** | **Breadth** | **Mean depth** | **Gini** |
| *Wolbachia* (wMel-like, supergroup A) | **12** | 0.688–0.788 | 7.1–45.2× | 0.383–0.455 |
| *Rickettsia* (*R. bellii*-like) | 8 | 0.610–0.624 | 9.7–63.7× | 0.412–0.510 |

**Supplemental Note S1: on the taxonomy of *Aphrophora sp.***

To set the identification of the vineyard *Aphrophora* in perspective, a brief review of the tangled taxonomic background of *A. permutata* is useful. Uhler (1876) described *Aphrophora permutata* following a preliminary listing of the name without description (Uhler 1872). He based his description on specimens from Utah, Colorado and, importantly for this work, “from near San Francisco.” In the only published revision of the genus Aphrophora, Doering (1941) separated and described as new two similar species, *A. fulva* and *A. maculosa*, based on subtle differences in the male subgenital plates and adjacent ninth abdominal sternites. She also asserted that these species, both less common than *A. permutata* and both described from type localities in California, could be distinguished by their lighter “tawny” color. In a subsequent PhD dissertation, Moore (1956) performed a comprehensive study of variation in male genital morphology of putative specimens of all three species. While he found considerable variation, he found no consistent relationship between genital morphology and nominal species designations and concluded that “the forms designated as the species *fulva* and *maculosa* represent types within the normal range of individual variation of *permutata*.” He proposed synonymizing *A. fulva* and *A. maculosa* with *A. permutata*.

Moore’s thesis work was never published. As a result, the species have not been formally synonymized. This has direct bearing on the work reported here because the identity of the taxon we report as *A. permutata* has been contested. Kelson (1964) published the only *A. permutata* life history study, based on a population in the Berkeley-Oakland Hills. We sampled our reference *A. permutata* population from the same locale. Hamilton (1982) identifies the species Kelson studied as *A. fulva*, based on the assertion that *A. fulva* is a pine-feeding species and that *A. permutata*’s primary host is Douglas fir (*Pseudotsuga menziesii*). *Aphrophora fulva* was described from specimens collected on *Pinus sabiniana* (Grey pine) in the Sierra Nevada mountains near Kernville (Doering 1941), in an area apparently just outside the natural range of Douglas fir (The Gymnosperm Database, 2026). *Aphrophora maculosa* was described without host information from the San Jacinto Mountains near Palm Springs, California in an area where Big cone Douglas fir, *Pseudotsuga macrocarpa*, a close relative of Douglas fir occurs mixed with Coulter pines (*Pinus coulteri*) (The Gymnosperm Database, 2026).

We believe Kelson’s original identification of the Berkeley-Oakland Hills taxon as *A. permutata* is correct: 1) Doering herself determined Kelson’s samples to be *A. permutata* (Kelson 1964). 2) Our results demonstrate that the taxon we studied, including samples from the Berkeley-Oakland Hills, lives as adults on both pines and Douglas fir. This calls into question Hamilton’s distinction between a “Western pine spittlebug” identified as a *A. fulva* and a “Douglas fir spittlebug” identified as *A. permutata*. 3) Dissection of male genitalia showed that our specimens, while exhibiting some of the variation Moore (1956) observed, best fit Doering’s illustration of *A. permutata*. On this basis we identify the taxon Kelson studied as *A. permutata*.

**Supplemental Methods 1: Modeling sequencing depth against inferred population assignment**

We modeled sequencing depth, as estimated with samtools v1.19.2 (Danecek et al., 2021) against the cluster assignments of each individual in the autosomal PCA, with a Kruskal–Wallis test, a one-way ANOVA, and pairwise contrasts via Mann–Whitney tests with Benjamini–Hochberg (Benjamini & Hochberg, 1995) correction. Effect size was quantified as epsilon-squared, the rank-based analogue of eta-squared (Tomczak & Tomczak-Łukaszewska, 2014). We also correlated depth against the first three principal components, ancestry components from admixture analyses at K=2 and K=3, indexing assignment strength of ancestral population assignments, and the Shannon entropy of the ancestry vector, which indexes how admixed an individual appears, with Spearman rank correlations (ρ). To assess significance, we implemented t-approximation t = ρ√((n − 2)/(1 − ρ²)) on n − 2 = 45 degrees of freedom. We validated the approximation against a permutation null of 100,000 shuffles; the two agreed to within 0.007 across all tests, so the approximation was retained. We then corrected all twelve correlation tests together with the Benjamini–Hochberg false discovery rate procedure. As the two Pine Mountain (PM) individuals carried roughly 90% of the (non-significant) depth–apparent-admixture association, were the two most admixed-looking individuals in the dataset, and had the lowest coverage, we replicated the nuclear PCA and admixture analyses without these two PM individuals to test for any effect they might have on the inferred population structure.

**Supplemental Methods 2: Association of endosymbiont infection with locality and climate**

We utilized the Open-Meteo archive API to retrieve daily average, peak, and low air temperatures at a 2m height for all localities between 1995 and 2024. These three decades of climate data allowed us to calculate ten specific thermal indices (Table S6).

We implemented statistical evaluations of endosymbiont distribution within Python using the NumPy (Harris et al., 2020), SciPy (Virtanen et al., 2020), pandas (McKinney, 2010), and GeoPandas (Fleischmann, 2026) libraries. Recognizing that individuals within a given locale lack independence, we emphasize site-level assessments as our primary metric and regard individual-level data as descriptive. We utilized Fisher's exact tests to evaluate site-level correlations between endosymbiont infection and binary variables. To compare continuous environmental metrics—such as elevation and thermal indices—between sites with and without infection, we employed Mann–Whitney U tests. We further assessed monotonic relationships between infection prevalence and site characteristics using Spearman's ρ and Kendall's τ across the nine sampling localities.

We examined the spatial structure of endosymbiont infection via permutation, shuffling infection status across 20,000 iterations to compare the observed variance among sites against a null distribution. Spatial aggregation was further tested by contrasting the mean great-circle distance between infected localities against 20,000 random site triplets. To determine the endemism of nuclear ancestry, we permuted cluster labels across individuals, assessing whether the maximum cluster count observed at any single site exceeded null expectations. To disentangle the influence of mitochondrial lineage from geographic origin, we repeated the site-structuring permutation specifically within the mtC mitochondrial lineage (n = 23). All statistical tests were two-sided. Given the exploratory nature of these analyses, we did not adjust for multiple comparisons.
